# An essential dynamics-based elastic network model to unravel the conformational dynamics of DNA, RNA, and protein-nucleic acid complexes

**DOI:** 10.64898/2026.03.11.710985

**Authors:** Marco Cannariato, Domenico Scaramozzino, Byung Ho Lee, Marco A. Deriu, Laura Orellana

## Abstract

The flexibility of DNA and RNA is known to play a central role in numerous biological processes, including chromatin organization and gene regulation. While a wide range of computational approaches have been developed to investigate the conformational dynamics and flexibility of proteins, analogous methods for nucleic acids remain comparatively underexplored. Elastic Network Models (ENMs) – coarse-grained mechanical representations in which macromolecules are modeled as networks of nodes connected by elastic springs – have been successfully applied to proteins, often allowing to capture experimentally observed conformational changes through a small number of harmonic normal modes. Building on a previously validated three-bead ENM for RNA, here we introduce edENM, an essential dynamics-refined ENM for DNA, RNA, and protein-nucleic acid complexes, parametrized using a diverse set of Molecular Dynamics simulations. The vibrational modes of the new edENM show good agreement with NMR data and experimental ensembles, while avoiding the unrealistic and localized deformability of previous ENM parametrizations. Additionally, we integrated this new edENM into eBDIMS, a Brownian Dynamics-based framework that enables the simulation of large-scale and anharmonic conformational transitions in protein assemblies. In this way, we are now able to explore functional motions in large protein-nucleic acid complexes such as chromatin subunits and ribosomes.

## Introduction

A key characteristic of nucleic acids is their dynamic and flexible nature^1^, which is important for their biological function. DNA flexibility is fundamental to reach a proper packing of the genetic material within the cell nucleus^2–5^ and RNA dynamics has gained growing interest due to the recent discoveries in its structure-to-function relationships^6–9^. Large-scale conformational events can be crucial to achieve specific functions^10^, as well as in complex phenomena like chromatin reorganization^2^. The dynamics and flexibility of nucleic acids are deeply influenced from their interactions with proteins^11^. Protein conformational changes are pivotal for protein function^12^ and that these are known to be conserved along evolutionary processes^13,14^. Protein flexibility is in turn largely influenced by interacting partners since local changes can propagate through the structure of the macromolecular complex and alter its overall dynamics^15^. Thus, a proper comprehension of the molecular features characterizing protein-nucleic acid interactions influenced by short time-scale phenomena^16^ is necessary to understand the functional dynamics of these protein-nucleic acid complexes.

Among the techniques that have been employed to investigate the flexibility of biomacromolecules, the gold standard is Molecular Dynamics (MD)^17^. MD allows to simulate with atomistic details the dynamics of proteins and nucleic acids, while reproducing experimental measures and assisting in their interpretation^18–20^. MD simulations have provided pivotal insights into a variety of biological processes^17,18^. Principal component analysis (PCA) of MD simulations, also referred to as essential dynamics (ED)^21^, has also offered detailed insights into molecular flexibility^22,23^, capturing large-scale and biologically relevant motions^24^. Despite these advantages, MD simulations face major challenges, such as the significant computational demand and the so-called “sampling problem”^12^. Several coarse-grained (CG) approaches have been developed to overcome the computational challenges posed by atomistic MD^25,26^. In the CG representation, the molecular structure is simplified in order to retain only the physical/chemical properties of major interest, reaching a compromise between computational efficiency and simulation accuracy.

Among CG methods, Elastic Network Models (ENMs) describe the structure of macromolecules as a network of elastic springs in the harmonic regime^27–29^. Despite the simplification of the free energy landscape with the harmonic potential of ENMs, these models have proven to be effective in describing the direction of biologically relevant motions, such as cooperative domain movements^23,30–35^. Yet, ENMs can often struggle to capture the anharmonicity of large-scale motions due to their intrinsic near-equilibrium small-oscillation approximation, which has made the application of ENMs more difficult to nucleic acids, that are typically assumed to undergo more pronounced anharmonic motions. In proteins, ENMs often use a single node per amino acid localized at the position of the C^α^ atom^29^ and the criteria to define the network connectivity and the spring constants differ significantly^36^. The few ENMs proposed for nucleic acids differ for the number and localization of CG nodes per nucleotide, as well as for the criteria to define the spring constants accounting for the bead-bead interactions^37–40^. No consensus exists for ENMs of protein-nucleic acid complexes^41–43^, as the main challenge has been lying in the lack of reliable experimental data to validate any ENM force-field.

A step forward in the definition of an optimized ENM for proteins made use of extensive MD simulations as benchmark for the ENM parametrization instead of the traditionally used B-factors^31^. As a matter of fact, atomic fluctuations from crystallographic experiments are strongly influenced by crystal packing and refinement errors^44,45^, which can alter the fluctuations from the intrinsic dynamics. On the other hand, MD data provide a description of both short- and long-time motions, also accounting for anharmonicity effects. The ED-based ENM (edENM) for proteins, refined on MD data and developed in Orellana et al.^31^, was shown to outperform standard ENM approaches both regarding global and local flexibility. The edENM force-field for proteins was also coupled to a biased Brownian dynamics (BD) simulation framework^46^ to obtain a transition path-sampling algorithm, eBDIMS^47^, which allows to explore non-linear transition pathways between end-state conformations and generate reliable protein intermediates^48^. Recently, eBDIMS has been improved to generate transition intermediates for larger assemblies from cryogenic electron microscopy (cryoEM)^49^.

In the context of nucleic acids, there have been a few approaches to identify best-performing ENM topologies, using experimental structures, MD, and SHAPE experiments^37,40^. These studies have been often limited to either RNA or DNA, and they have generally proposed two distinct parametrizations for the two nucleic acids. In these studies, a three-bead model has been shown to provide the best compromise between accuracy and simplicity^40^. While different formulations of spring constants were proposed, these were found to correlate differently to experimental data^37^ and, to the best of our knowledge, no refinement of nucleic-acid ENM has ever been attempted based on MD data. For this reason, we have performed a systematic refinement of the classical three-bead ENM model for DNA and RNA, making use of extensive structural-dynamical data from MD simulations and structural ensembles from the Protein Data Bank (PDB)^50^. This new edENM force-field expands the previous edENM model developed for proteins in Orellana et al.^31^, allowing to employ a single ENM parametrization for both RNA and DNA, and improves the correlation between the computed and experimental conformational changes with respect to traditional ENM parametrizations, generating more collective harmonic motions. We have also integrated the new edENM force-field into eBDIMS^48,49^, so that we are now able to explore large-scale conformational changes in DNA, RNA, and large protein-nucleic acid complexes.

## Materials and Methods

### Parametrization workflow

The parametrization of the new edENM (Fig. 1) has been performed in two phases: (1) parametrization of the ENM for DNA and RNA; (2) definition of the interface contacts between the nucleic acid and the protein, and integration of the nucleic acid edENM with the protein edENM developed in Orellana et al.^31^. For the topology of the nucleic acid edENM, we started from the three-bead sugar-base-phosphate (SBP) model proposed by Pinamonti et al.^40^ (Supplementary Fig. 1). We chose the SBP topology as it was considered to be a good compromise between network complexity and model accuracy^40^. In the original SBP model, all springs have the same force constant, a spatial interaction cutoff of 9 Å, and each nucleotide group is represented by a single bead, C1’ representing the sugar, C2 the nitrogenous base, and P the phosphate group (Supplementary Fig. 1)^40^.

**Figure 1.**
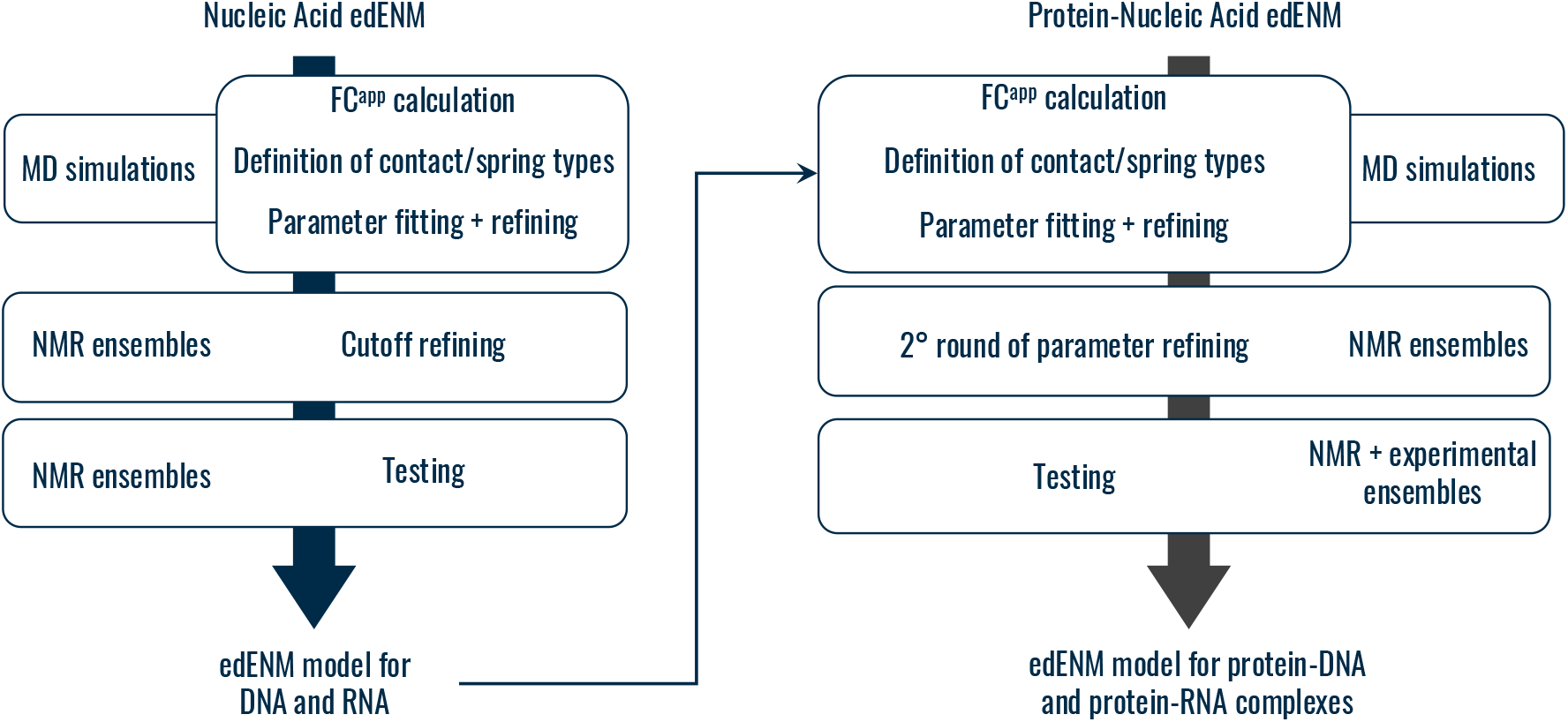
Workflow for the parametrization of the new edENM for (left) nucleic acids and (right) protein-nucleic acid complexes.

After defining the initial SBP topology, we used MD-based parametrization coupled with further refinement based on experimental data, to develop the new edENM for DNA, RNA, and protein-nucleic acid complexes. In the first phase of the parametrization for DNA and RNA, we estimated the apparent force constants between all P, C1’, and C2 atoms from a training set of nucleic acid MD trajectories (see below), as^31^:

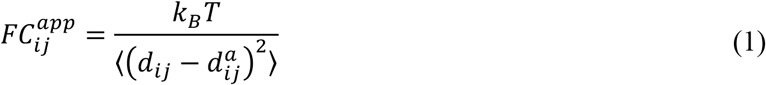

where *d*_*ij*_ represents the instantaneous distance between atoms *i* and *j*, 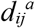 the average value of this distance along the trajectory, *k*_*B*_ the Boltzmann constant, *T* is the temperature, and ⟨…⟩ stands for the averaging across all MD frames. To obtain a first description of the edENM spring constants, the values of 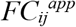 were mapped against the corresponding average distances 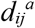, and the 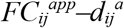 relationships were used to define different contact (spring) types. When a distance-dependent relationship was observed, this was fitted using an inverse exponential function as in Orellana et al.^31^:

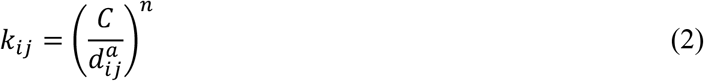

where *k*_*ij*_ represents the spring force constant in the edENM, and *C* and *n* are the parameters associated with the strength and the distance-dependent decay of the spring constant, respectively. A first estimation of these parameters was performed by minimizing the standard error of the *k*_*ij*_ regression curve against 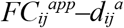 MD values. These parameters were then refined performing a random search in the neighborhood of the estimated variables, using a random combination of 1,500 values. Optimal values of *C* and *n* were subsequently found by comparing the vector alignment (overlap) between the normal modes (NMs) from Normal Mode Analysis (NMA) of the edENM and the ED eigenvectors from each MD trajectory (see below). After this refinement stage, the best parameters that maximize Root Mean Square Inner Products (*RMSIPs*, see below) were compared using different cutoff values, ranging from 9 to 14 Å. In this step, the NMs were also compared against Principal Components (PCs) derived from experimental ensembles of DNA and RNA molecules obtained from multi-model PDB files, such as those obtained from solution Nuclear Magnetic Resonance (NMR). Once the optimal parameters were selected, we tested the resulting edENM against the remaining MD simulations and an additional testing dataset of NMR and multi-state ensembles (see below). The performance of yhis optimized edENM was compared against the uniform-spring SBP-ENM from Pinamonti et al.^40^.

In the second parametrization phase, we estimated the apparent force constants 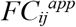 for protein-nucleic acid interface contacts from an additional set of training MD simulations that include protein-nucleic acid complexes. A first optimization of spring constants was performed making using of the inverse exponential function from Eq. (2) every time a distance-dependent relationship was observed. A random search in the neighborhood of the *C* and *n* parameters was performed using a combination of 1,100 values and considering several interaction cutoffs. The parameters of the edENM spring constants and the cutoff for protein-nucleic acids contacts were further refined using the training MD trajectories and performing an additional refinement based on experimental ensembles from NMR. Finally, the performance of the model was tested considering a separate set of MD trajectories, the NMR dataset, as well as a dataset of structural ensembles that characterize conformational changes in protein-nucleic acid complexes (see below).

### Dataset of Molecular Dynamics (MD) simulations

To perform an ED-based parametrization, a set of MD simulations of different systems is necessary, possibly simulated with different force-fields and in different conditions. We retrieved several MD trajectories from the existing literature using *MDverse*^51^, and making use of *Zenodo, OSF*, and *figshare* online repositories. Molecular systems composed of DNA, RNA, as well as protein-nucleic acid complexes were considered. In total, we retrieved separate MD simulations for 11 DNA, 5 RNA, and 10 protein-nucleic acid systems. For each system, the number of available replicas ranged between 1 and 20, with simulation lengths ranging from a minimum of 30 ns to a maximum of 2 μs. Detailed information about the MD dataset is reported in Supplementary Table 1. For nucleic acids, 12 simulations were included for training and 4 for testing, while 7 simulations of protein-nucleic acid complexes were included for training and the remaining 3 for testing. For each simulation, we discarded the first 10 ns as these generally include initial equilibrations. Also, large-scale and anharmonic conformational changes might arise in longer simulations, and these motions are generally difficult to capture with NMs from ENMs due to their intrinsic harmonic and small-amplitude nature. For this reason, we split each MD trajectory in smaller windows. For total simulation lengths < 1 μs, we extracted windows of 50 ns, while for longer simulations we used windows of 100 ns. Since some local conformational changes might end up being split into adjacent windows, we also considered an overlap of 10 ns between adjacent windows. In cases where the total simulation length was < 50 ns, the whole trajectory was taken as a single window.

On the resulting “windowed” trajectories, we performed Principal Component Analysis (PCA). PCA is a statistical technique used to capture the low-dimensional variables that explain the majority of the variance within high-dimensional data. PCA of MD simulations is usually referred to as Essential Dynamics (ED) and was shown to provide meaningful insights into protein dynamics and flexibilities^21,23^. The input of PCA is an *n* × *3N* coordinate matrix, ***X***, where *n* is the number of conformations in the MD trajectory and *N* the number of atoms-residues, usually considering only C^α^ atoms for proteins and P atoms for nucleic acids. From ***X***, the elements of the symmetric covariance matrix, ***C***, can be calculated as:

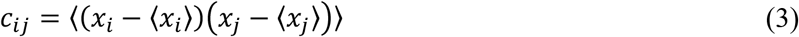

where the brackets ⟨…⟩ indicate the average over all MD simulation frames. Eigenvalue-eigenvector decomposition is then used to diagonalize the covariance matrix as:

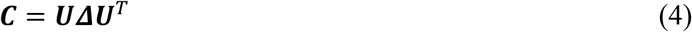

where the diagonal matrix ***Δ*** contains the eigenvalues of ***C***, while the matrix ***U*** contains its eigenvectors, representing the Principal Components (PCs). Eigenvalues are sorted in descending order and are directly proportional to the amount of variance explained by the corresponding PC eigenvector. Each PC eigenvector can be directly compared to NMs via overlaps and *RMSIP* scores (see below).

### Dataset of experimental ensembles and conformational changes

To refine the model parameters so that the dynamics of the edENM also captures experimentally observed conformational changes, we also made use of experimentally available nucleic acid structures and protein-nucleic acid complexes from the PDB^50^. For systems composed of solely nucleic acids (DNA or RNA), we retrieved a total of 29 ensembles, 25 of which are single NMR models, 3 are ensembles of multiple NMRs or multi-model cryoEM structures, and one is an ensemble of distinct models from cryoEM. Out of these 29 ensembles, 10 were used for cutoff refining, and 19 for testing. The set of all PDB models is listed in Supplementary Table 2.

We also retrieved experimental structures of protein-nucleic acid complexes from NMR models. After removing duplicates and NMRs with only one single model uploaded, we retrieved a list of 208 potentially usable NMR structures. Then we inspected these models and retained only those with a relevant conformational diversity. This was enforced by computing Root Mean Square Fluctuations (*RMSFs*) and Root Mean Square Deviations (*RMSDs*). First, we retained only those protein-nucleic acid complexes where maximum *RMSD* values between any two models in the NMR ensemble were > 3Å. Then, we removed portions of the structure where *RMSFs* were localized at the N- and C-termini of protein chains, which is common in NMR ensembles. This was done since these types of localized conformational changes at the protein tails are not the typical targets of ENM analyses, that tend to provide meaningful information on more collective motions. Finally, we re-computed *RMSD* values on the models without the flexible tails and retained only those with a global *RMSD* still > 3Å. These *RMSD*- and *RMSF*-based filters led us to retain a total of 21 significant protein-nucleic acid NMRs (Supplementary Table 3), that were used for edENM parameter refinement and testing.

In the case of ensembles and conformational changes in protein-nucleic acid complexes coming from sets of different X-ray/cryoEM structures, we carried out a search of ensembles by performing a screening of the PDB and UniProt database^52^. First, all proteins with solved 3D structures and reviewed UniProt entries were considered (35,348 entries, as of May 2024). Then, we retained only those PDBs where at least one protein entity and one nucleic acid entity were present. UniProt IDs having only one associated PDB were discarded, reducing the suitable entries to 4,258. UniProt IDs sharing the same PDBs, e.g., due also to the presence of heteromeric protein-protein complexes, were merged into the same UniProt-PDB combination, which led us to 2,374 unique groups of PDB models. Then, entries with one or more proteins having a molecular weight higher than 400 kDa were filtered out in order not to consider gigantic protein-nucleic acid assemblies for NM testing. Then, we computed *RMSDs* between all PDBs via *US-Align*^53^, maintaining only those with *RMSDs* > 3 Å in order to ignore ensembles with negligible conformational diversity. Finally, we filtered out structures with low-resolution (> 5Å) and removed assemblies including more than 10 different polymer entities, which led to a final list of 265 potential ensembles/conformational changes. This final list of ensembles was manually inspected to exclude inadequate cases due to a variety of reasons, e.g., nucleic acids with sequences not matching in the different conformers, ensembles resulting in disconnected regions due to many missing residues, high *RMSDs* solely due to the motion of terminal tails, etc. Eventually, the final testing set was composed of 26 structural ensembles with at least 2 distinct PDB models that satisfied all our filtering criteria. The full list of ensembles with the included PDB models is reported in Supplementary Table 4. For each set of testing NMR models and conformational ensemble we carried out PCA and extracted the experimental PCs that explain the majority of the variance within the ensemble. PC eigenvectors were then compared with NMs from edENM through overlap and *RMSIP* scores (see below).

### Normal Mode Analysis (NMA) and comparison metrics for edENM optimization and testing

Normal Mode Analysis (NMA) is a mathematical technique used to extract the vibrational features of a structure. In the past decades, NMA has been widely applied to investigate the dynamics of proteins, nucleic acids, and biomacromolecules in general^27,54^. NMA relies on the definition of the Hessian matrix, that contains the second derivatives of the potential energy of the system. Atomistic NMA often requires a preliminary minimization of the system and employs MD-based force-fields accounting for energy variations in bond lengths, bond angles, and dihedral angles, as well as long-range interactions^54^. On the contrary, ENM-NMA relies on a simpler harmonic potential that treats the structure as a network of elastic springs, and allows to avoid the minimization step as it relies on the assumption that the ENM conformation is already in the energy minimum^27,29^. To compute NMs for comparison with ED eigenvectors from MD trajectories, the ENM was built starting from the coordinates of the MD frame closest to the average conformation within each MD window. Experimental PDBs were instead used directly as reference topologies when comparing NMs to experimental PCs.

Once the molecule topology is determined and spring parameters are established, the 3*N* x 3*N* edENM Hessian matrix ***H*** is assembled as a combination of 3 x 3 symmetric sub-blocks 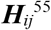. Each sub-block contains the stiffness information about the interaction between nodes *i* and *j*:

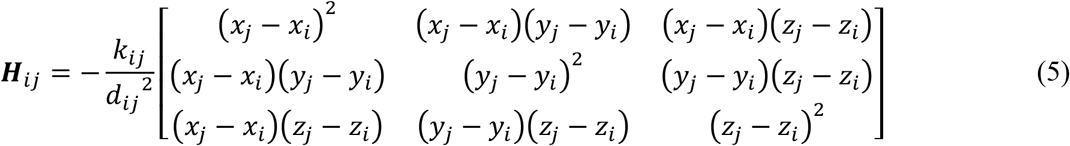

where *k*_*ij*_ is the stiffness of spring in the network connecting nodes *i* and *j, d*_*ij*_ their distance, and *x, y, z* are the 3D coordinates derived from the X-ray, NMR, cryoEM model or the MD simulation frame. Once the Hessian matrix is assembled, its eigenvalues and eigenvectors are computed based on a standard matrix diagonalization, as already reported in Eq. (4) for the PCA covariance matrix ***C***. In this case, the diagonal matrix ***Δ*** contains the eigenvalues of ***H***, ordered by ascending value, and the column of ***U*** provide the full set of 3*N* eigenvectors. Each eigenvalue-eigenvector pair is associated to a specific normal mode (NM) of vibration: the eigenvalues describe the NM vibrational frequencies and are also associated to the NM energy and amplitude (low-frequency NMs are also the lowest-energy and generally more collective ones), while the eigenvectors describe the 3D directions along which NMs take place. The first 6 NMs describe zero-frequency motions associated with global translations and rotations and are discarded from following analysis.

The effectiveness of the method in capturing conformational changes was assessed by comparing the directions of NMs from the edENM to: (1) the ED vectors from MD, (2) the PCs from NMR and multi-model PDB structures, and (3) the PCs from a collection of different PDB structures in a common experimental ensemble. For this purpose, we computed overlaps between NMs and PCs. The overlap *O*_*ij*_ between a NM eigenvector ***ν***_*i*_ and a PC eigenvector ***w***_*j*_ from an ensemble is computed according to the following equation:

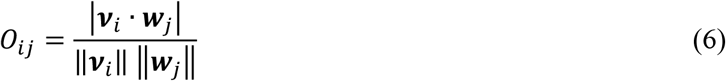

where ‖***u***‖ indicates the Euclidean norm of the vector ***u***. An overlap of 0 implies that the two vectors are orthogonal, while for two matching (parallel) vectors the overlap is 1. On the other hand, the *RMSIP* expresses the similarity between two larger vectorial sub-spaces described by *n* vectors ***ν*** and ***w***, and it was computed as:

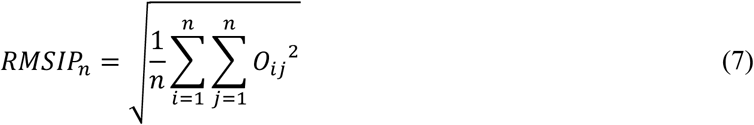

Typically, *RMSIP* calculations are restricted to the first NMs, e.g., *n* = 3, 5, or 10 modes, as the lowest-frequency motions typically represent functional motions^14,30,35^. Another useful metric to assess the global vs. localized nature of conformational motions from PC or NM eigenvectors is the collectivity degree *κ*, which is defined as^30^:

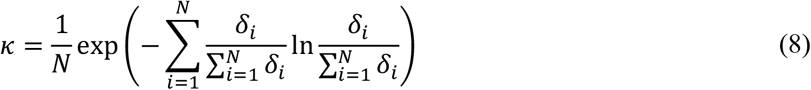

where *δ*_*i*_ stands for the absolute value of the displacement in the PC or NM eigenvector at residue *i*. The collectivity degree varies from a theorical minimum of 1/*N*, if only one residue participates to the motion, to a maximum of 1, if all residues are uniformly involved in the structural change. The former case is associated with extremely localized motions, the latter with highly collective vibrations.

### Implementation in eBDIMS for path-sampling of conformational transitions

After assessing the performance of the new edENM for nucleic acids and protein-nucleic complexes, we integrated this new force-field into the eBDIMS^47,48^. eBDIMS is a CG path-sampling algorithm that relies on a protein ENM representation, a BD simulation framework, and can be used to track intermediate states along transition pathways by using a Dynamic IMportant Sampling (DIMS) bias^48^. eBDIMS was shown to produce realistic protein intermediates along transition pathways and capture experimentally observed intermediates. eBDIMS was also recently upgraded to a more efficient version, eBDIMS2, that can now simulate large-scale and complex transitions in gigantic (MDa) protein assemblies from cryoEM^49^. We implemented the new edENM parameters for nucleic acids and protein-nucleic acid contacts into eBDIMS2, so that we can now simulate large-scale conformational transitions not only in proteins, but also in DNA, RNA, as well as a wide variety of protein-nucleic acid systems, with virtually no restriction on system size, macromolecular architecture, and motion complexity. Here, we applied eBDIMS to simulate conformational changes in four different systems: a classical hinge-bending motion in the *BtCoV-HKU5* 5’ proximal stem-loop 5 RNA (∼40 kDa), a localized RNA transition from the central-duplex to the bent-duplex conformation of *A. Thaliana* Argonaute 10 (∼120 kDa), a large-scale opening of an *H. Sapiens* telomeric trinucleosome (∼560 kDa), as well as a hinge-bending motion in the large 18S rRNA of an *H. Sapiens* pre-40S ribosomal subunit (∼1.3 MDa).

## Results

### Parametrization and testing of the nucleic acid edENM

The distribution of 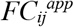 values as a function of the inter-bead distances for P, C1’, and C2 atoms were computed from the training set of MD trajectories as described in the Methods section. Putative differences between DNA-only (Supplementary Fig. 2) and RNA-only (Supplementary Fig. 3) systems were investigated, but no major distinction was found to justify a different parametrization. Therefore, for the sake of simplicity, one edENM force-field was parametrized for both DNA and RNA systems. The overall distribution of 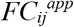 values in the training trajectories is shown in Fig. 2, which made us define four distinct interaction types.

**Figure 2.**
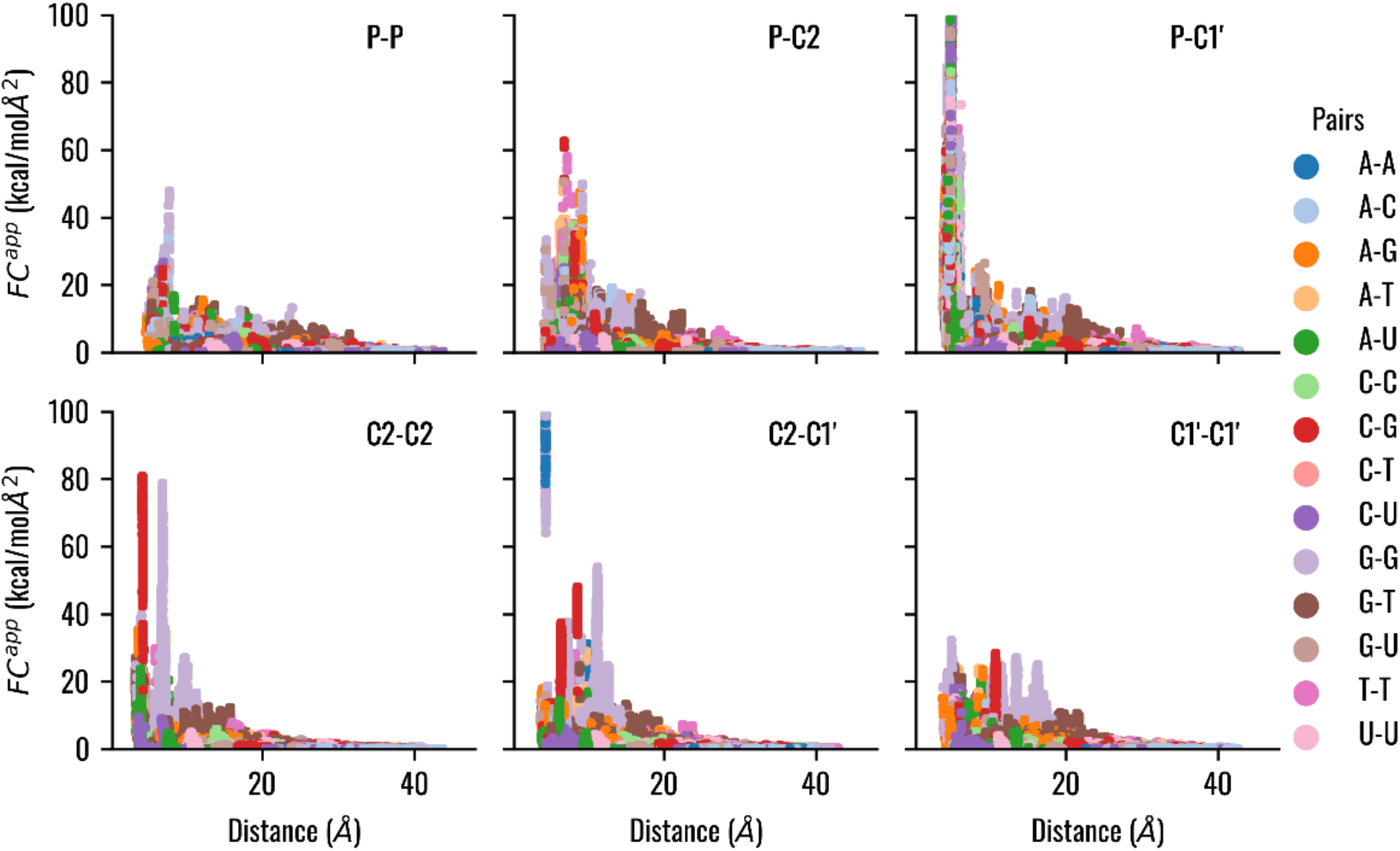
Distribution of the apparent force constants 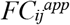 as a function of the average distance 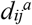 in the training MD dataset of nucleic acids systems, aggregating both DNA- and RNA-only systems. The apparent force constants between different interacting couples are shown and the different nucleotide- nucleotide interactions are highlighted with different colors.

The first type of interaction is the “*covalent*” interaction, defined between C2 and C1’ beads within the same nucleotide. This interaction represents the strong bond linking the sugar ring and the base and contributes the structural stability of the nucleoside. The 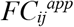 for this interaction has values over 150 kcal/molÅ^2^, reaching peaks even at 800 kcal/molÅ^2^ (Supplementary Figs. 4-5). Depending on the type of bases, such peaks were localized at two specific distances between C2 and C1’ beads: the higher one with purine and the lower one with pyrimidine (Supplementary Fig. 4). For these “*covalent*” interactions, we looked for parameters that lead to distance-independent stiffness values, i.e., with *n =* 0 from Eq. (2). The second type of interaction is the “*pseudo-covalent*” interaction defined between P and C1’ beads within the same nucleotide, forming the backbone of the nucleic acid. The 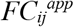 distribution for these bead pairs was quantitatively different from all remaining ones. In fact, we observed high 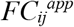 values, up to ∼150 kcal/molÅ^2^, for consecutive P-C1’ couples comprising of the nucleic acid backbone. For this “*pseudo-covalent*” interaction, we looked for optimal *C* and *n* parameters that allow to follow a distance-dependent law according to Eq. (2). The third recognized interaction type is the “*Hbond*” interaction for close C2-C2 pairs. This interaction represents the hydrogen bond-driven base-pairing. We observed two distinct peaks for this interaction (Fig. 2). The first peak was found to correspond to classic base-pairing interactions in the distance range 4.1-4.4Å, and the second to G-G contacts in the range 6.7-7.0Å. This surprising G-G interaction was related to MD simulations containing a β-hairpin (Supplementary Fig. 6). We confirmed that these peaks are independent to sequential distances (Supplementary Fig. 7), and we modelled these interactions with a distance-independent spring law, only characterized by the parameter *C*. All remaining interactions were grouped in a fourth type of interaction, which was termed “*Van der Waals*” and modelled according to Eq. (2).

The range of *C* and *n* parameter values that was employed in the random search refinement for the four interaction types is reported in Supplementary Table 5. An example of fitting interval based on minimum and maximum values for the “*pseudo-covalent*” and “*Van der Waals*” distance-dependent interactions is shown in Supplementary Fig. 8. After refinement, the C1’-C2 “*covalent*” interactions were associated with constant values of 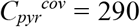 kcal/(molÅ^2^) for pyrimidines, and 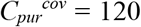 kcal/(molÅ^2^) for purines. The C2-C2 “*Hbond*” interactions were optimized with a constant value of *C*^*Hbond*^ = 65 kcal/(molÅ^2^). The interactions between the P and C1’ atoms in the “*pseudo-covalent*” interactions were optimized with a distance-dependent law, with *C*^*psc*^ = 20 and *n*^*psc*^ = 2.8. Finally, all remaining “*Van der Waals*” interactions were modeled with *C*^*VdW*^ = 25 and *n*^*VdW*^ = 1. At the end of the nucleic acid edENM parametrization, the optimal cutoff for “*pseudo-covalent*” and “*Van der Waals*” distance-dependent interactions was set to 11 Å. A visual representation of the resulting edENM topology is shown in Fig. 3, and compared to the traditional uniform-spring ENM^40^.

**Figure 3.**
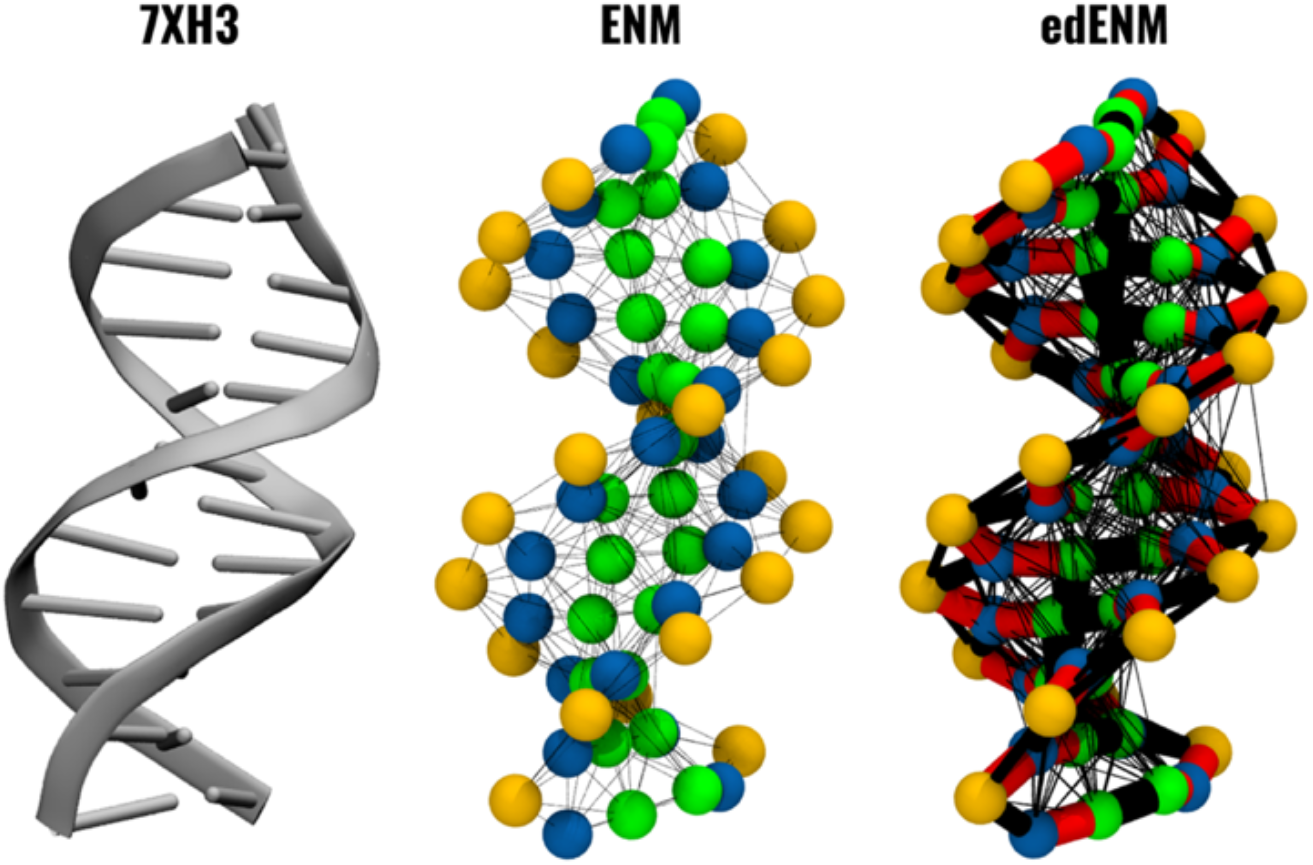
Elastic network topology for a double-stranded DNA helix (PDB ID: 7XH3). The standard uniform-spring SBP-ENM from Pinamonti et al.^40^ is compared to our refined edENM. P, C1’, and C2 atoms are represented as spheres, with P atoms in yellow, C1’ atoms in blue, and C2 atoms in green. Springs are represented with black cylinders with diameters proportional to the spring constant values. In the edENM, diameters have been saturated to a maximum value of 65 kcal/molÅ^2^, and spring constants exceeding this value (“*covalent*” and “*Hbond*” interactions) are represented as red cylinders.

After finding the optimal spring network connectivity and spring parameters, we compared our new edENM to the three-bead uniform-spring ENM proposed by Pinamonti et al.^40^. For sake of simplicity, the uniform-spring model from Pinamonti et al.^40^ will be referred to simply as ENM in the following. First, we compared the performance of the ENM and edENM in capturing the MD flexibility by comparing their NMs against the PCs of four different systems in the MD testing set (Supplementary Table 1). From this comparison, we observed that the proposed ED-based parametrization produces generally reasonable overlaps with MD simulation data and it slightly increases the *RMSIP* agreement between the lowest-frequency NMs and the trajectory PCs in the testing set compared to the uniform-spring parametrization (Supplementary Fig. 9).

Then, we compared the performance of the edENM in terms of its ability to capture apparent motions in the testing dataset of DNA- and RNA-only multi-model PDBs (Supplementary Table 2) and compared it to the ENM. Both ENM and edENM were generally able to describe the experimental PC motions via their low-frequency NMs (Supplementary Fig. 10, Fig. 4). Average values of *O*_*max*_ and *RMSIP* scores across the multi-model testing dataset were > 0.6 for both models (Supplementary Table 6), with maximum overlap scores reaching > 0.9, suggesting very good agreements between experimental and predicted motions. Moreover, most of the *O*_*max*_ scores were found between NM1 to PC1 (Supplementary Table 6), indicating that the lowest-frequency NMs from simplified ENM topologies have the ability to capture the main directions of motion from the experimental ensembles.

**Figure 4.**
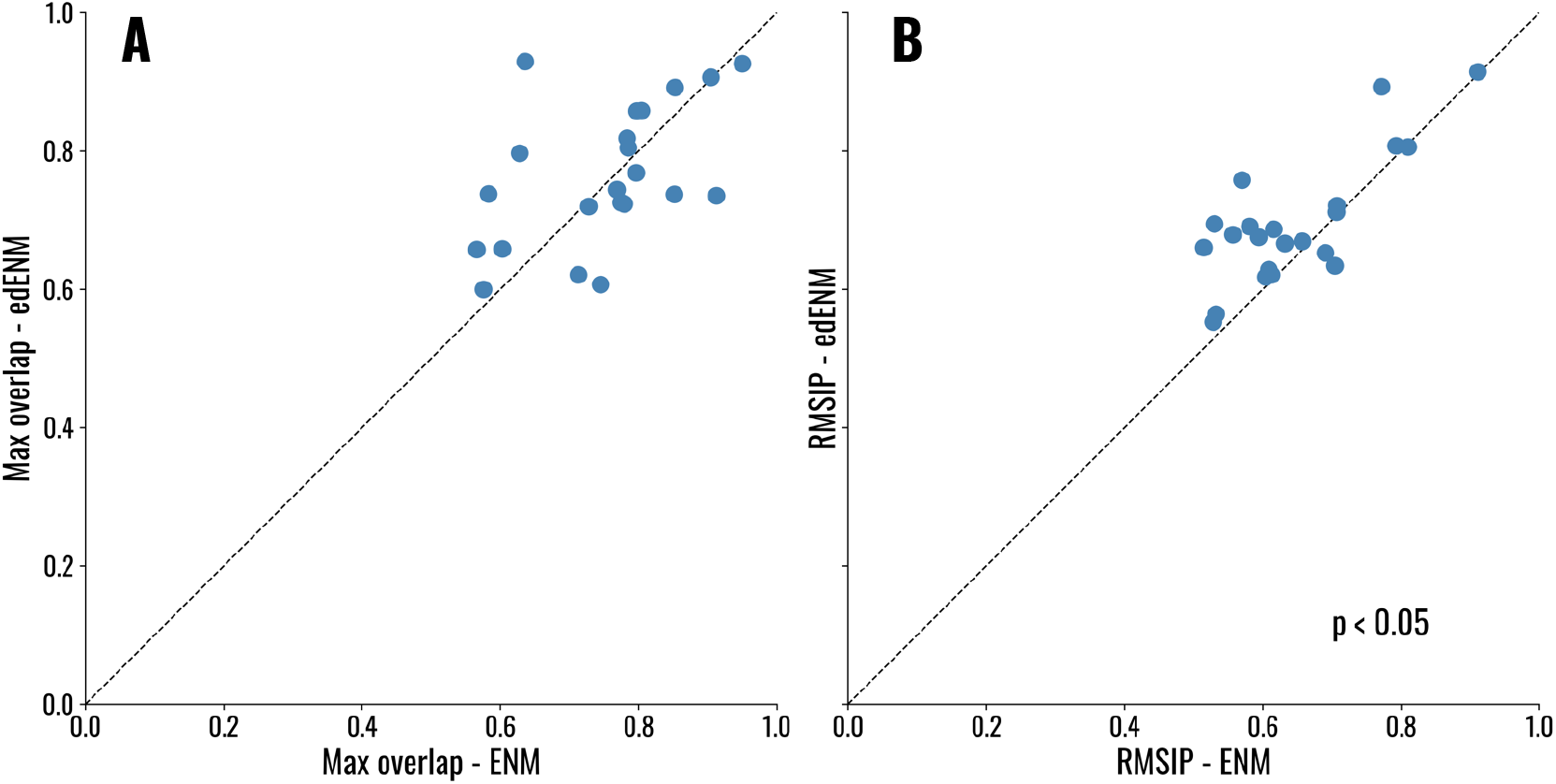
Comparison between the edENM and ENM in terms of ability to capture experimental PCs from the set of multi-model PDB structures of nucleic acids (Supplementary Table 2). (A) *O*_*max*_ for the set of first 3 NMs and experimental PCs. (B) *RMSIP* for the set of first 3 NMs and experimental PCs. *p*-values arising from a one-sided Wilcoxon test (edENM > ENM) between *O*_*max*_ and *RMSIP* scores are 0.33 and 4 × 10^−4^, respectively (Supplementary Table 6).

The performance of the edENM was satisfactory in terms of *O*_*max*_ scores, and in a few cases the ED-based parametrization provided a much higher overlap compared to the ENM. For instance, the opening-closing motion in the NMR of the U65 Box H/ACA small nucleolar RNA (PDB: 2PCV) was captured with 64% similarity by NM1 of the ENM, and with a remarkable 93% by the edENM (Fig. 5). We observed that the lower ENM similarity was due to an unrealistic transverse “breakage” of the RNA backbone in the dominant NM (Fig. 5E), which was not generated by the edENM due to its higher spring constants in the RNA backbone (see Fig. 3). Interestingly, in a few cases, *O*_*max*_ scores were slightly higher when using the simpler ENM (Supplementary Table 6). For instance, the opening-closing motion related to the experimental PC1 of a III-IV-V three-way junction of VS ribozyme (PDB: 2MTJ) was captured with 74% similarity by NM1 of the edENM, while exhibiting a higher overlap (85%) with NM2 of the ENM (Supplementary Fig. 11). However, looking in more detail at this case, we observed that the lowest-frequency mode (NM1) of the uniform-spring ENM gave rise to an extremely localized oscillation involving a detachment of the C2 bead in residue G47 (Supplementary Fig. 11), making this dominant NM completely uncorrelated with experimental motions from the NMR ensemble. As a result, the *RMSIP* of the three lowest-frequency modes was still higher in the optimized edENM (0.67) than in the ENM (0.59), as evident for most cases from Fig. 4B.

**Figure 5.**
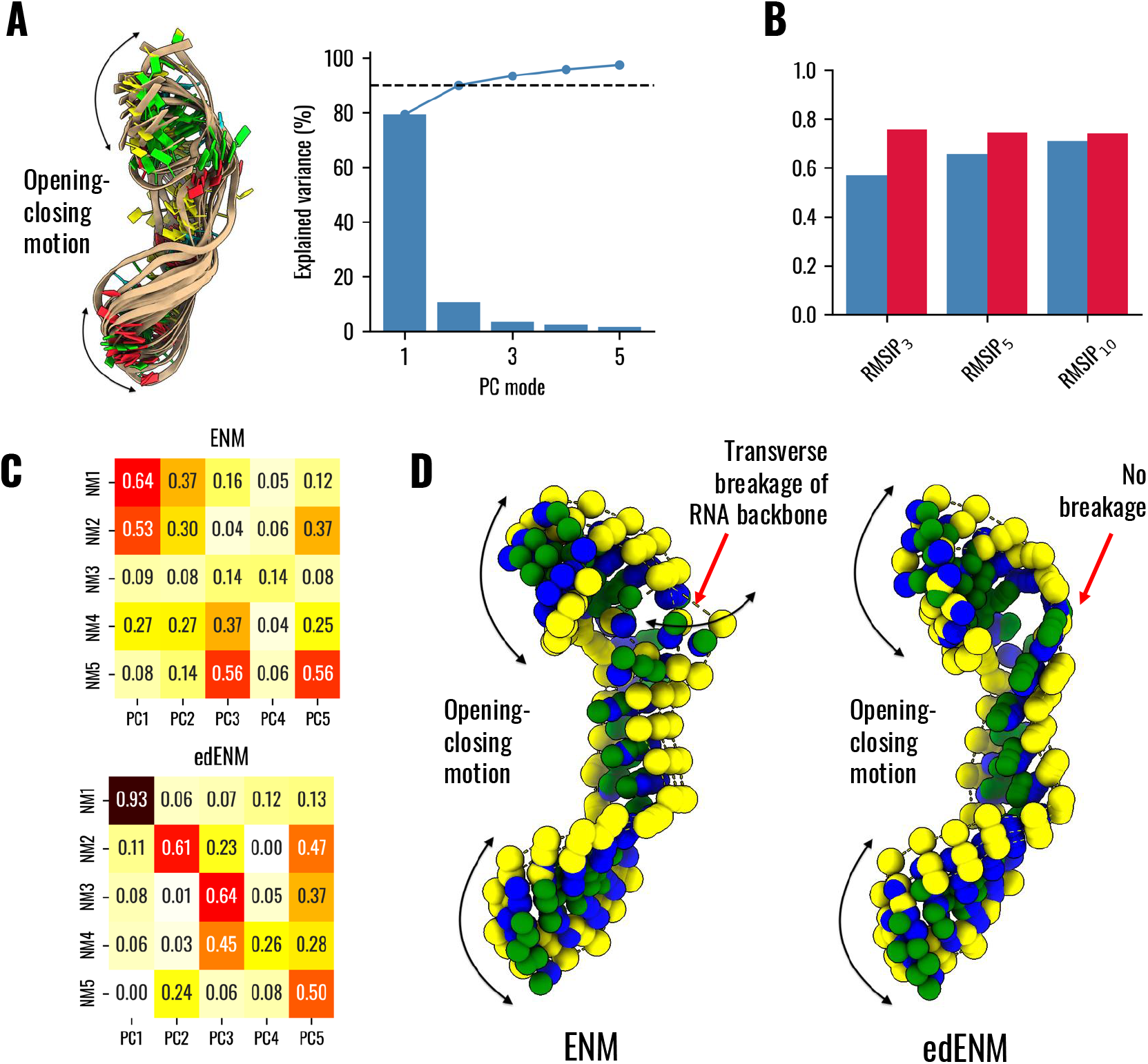
Comparison between ENM, edENM and PC modes from the NMR ensemble (PDB: 2PCV) of the U65 Box H/ACA small nucleolar RNA (snoRNA). (A) Ensemble of 20 models in the solution NMR ensemble used for PCA. The black arrows highlight the opening-closing motion apparent from the experimental ensemble, which dominates PC1. The graph shows the variance captured by each PC mode (blue bars), the cumulative variance of the first PC modes (continuous blue line), and a threshold of 90% variance (dashed black line). PC1 captures ∼80% variance and PC1+PC2 together allow to capture ∼90% variance. (B) Comparison between ENM (blue bars) and edENM (red) in terms of *RMSIP* scores (considering both 3, 5, and 10 modes) between NMs and experimental PCs. (C) Matrix of overlaps between the first 5 PC modes and the first 5 NMs from ENM and edENM. Maximum overlaps are found between NM1 and PC1 and reach 0.64 for ENM and an astonishing 0.93 for edENM. (D) Graphical representation of the motions associated with NM1 for ENM (left) and edENM (right). Beads represent P, C1’ and C2 positions across three deformed structures along the NM, with the same coloring as in Fig. 3. Note how both models allow to capture the opening-closing motion observed in the NMR ensemble, but the uniform-spring ENM shows a “breakage” (pronounced transverse oscillation) of the RNA backbone, which is prevented in the edENM due to stronger backbone interactions.

When the performance of the two models was evaluated across the entire dataset of multi-model structures, we found that *O*_*max*_ values in edENM were on average greater than in the ENM (Supplementary Table 6), even though the slightly positive difference (∼ +1.5%) was not found to be statistically significant (*p*-value ∼ 0.33). On the other hand, a clear significant difference was observed when comparing the lowest-frequency NM sub-spaces through *RMSIP* scores, with edENM significantly better performing (∼ +5%, *p*-value < 0.0005) than the uniform-spring parametrization (Supplementary Table 6). Taken together, these results suggest that, while correlations between individual NMs and experimental motions might not necessarily be higher in all cases, our ED-based model tends to generate more realistic harmonic motions, with no evident backbone breakage or extremely localized oscillations, and exhibits significantly higher values of sub-space similarity (*RMSIP*) with experimental PCs.

### Parametrization and testing of the protein-nucleic acid edENM

The distribution of protein-nucleic acid 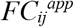 values as a function of the inter-bead distances was computed using the corresponding MD dataset for protein-nucleic acid systems in Supplementary Table 1. The relationship between distances 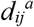 and 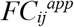 was inspected considering all possible combinations of contacts between different protein residues and nucleic acid bead types. Protein amino acids were divided according to their chemical nature, i.e., acidic, basic, polar, and hydrophobic. From the 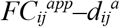 plots, we observed that the interactions between P atoms with polar or basic residues in proteins showed a marked distance-dependent decay with *n > 1*, while the remaining pairs had *n* closer to 1 (Supplementary Fig. 12). In case of C1’ beads, no interaction couple showed a clear decay with *n > 1* (Supplementary Fig. 13), while for C2 beads, we observed exponents greater than 1 in the case of acidic residues (Supplementary Fig. 14). Thus, we defined two types of interaction to describe the network of protein-nucleic acid contacts, i.e., “*strong*” and “*weak*” interactions, both following a distance-dependent decay law as for Equation (2). The former were defined for interactions between P beads with polar and basic residues, as well as interactions between C2 beads with acidic residues. The latter were used to describe all remaining connections between C^α^ atoms of the protein and nucleic acid beads. The parameters for “*strong*” and “*weak*” protein-nucleic acid interactions were fitted based on the MD-derived 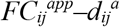 data. The range of parameters explored in the random search and in the subsequent refinement is reported in Supplementary Table 7. After refinement and optimization using NMR models of protein-nucleic acid complexes (Supplementary Table 3), we found optimal values *n*^*str*^ = 2.2, *C*^*str*^ = 25, and *C*^*weak*^ = 30 (*n*^*weak*^ = 1), with an optimal protein-nucleic acid interaction cutoff of 8 Å. Regarding the topology of the protein-nucleic acid interface, differently from the standard ENM approach, we introduced a limit in the number of springs connecting protein residues and nucleotides. For each pair of protein-nucleic acid residues, we considered only one elastic connection, specifically between the C^α^ atom of the protein and the closest bead in the nucleotide within the interaction cutoff (Supplementary Fig. 15A-B). This criterion was implemented to avoid excessive stiffening of the protein-nucleic acid interface.

We compared the NMA results arising from the protein-nucleic acid edENM to those obtained with a standard uniform-spring ENM, where all springs have equal force constants. To identify the contribution of the new nucleic acid parametrization, we also considered a “hybrid” model, that we call here edProt, which is a combination of the edENM force-field^31^ for the protein and the uniform-spring ENM parametrization of Pinamonti et al.^40^ for the nucleic acid (see Supplementary Fig. 15C). The performance of these three models (edENM, ENM, and edProt) was assessed in terms of their capability to capture ED modes from a test set of three MD simulations (Supplementary Table 1), experimental PCs from a set of NMR complexes (Supplementary Table 3), as well as experimental PCs from conformational ensembles (Supplementary Table 4). For two^56,57^ out of the three testing MD simulations, both edProt and edENM provided greater *RMSIP* values than ENM, with edENM performing generally slightly better than edProt (Supplementary Fig. 16), confirming the advantage of using an MD-refined parametrization for both the protein and the nucleic acid to better capture global motions of the system. Surprisingly, in one of the three simulations^58^, the edENM/edProt performance was lower. In this case, the trajectories focused on the movements of disordered histone tails in a nucleosome complex, which are hardly modelled by any ENMs, as confirmed by low values of *RMSIP* in all three models (< 0.4).

For the set of 21 NMR protein-nucleic acid complexes (Supplementary Table 3), the optimized edENM was found to provide higher similarity scores to the experimental PCs (*O*_*max*_: 0.59, *RMSIP*: 0.51) compared to the uniform-spring ENM (*O*_*max*_: 0.52, *RMSIP*: 0.45) and edProt (*O*_*max*_: 0.45, *RMSIP*: 0.49, Supplementary Table 8). The comparison between edENM and ENM showed significantly improved scores in edENM for both *O*_*max*_ and *RMSIP* (*p*-value < 0.05, Supplementary Table 9, Supplementary Fig. 17), confirming the higher performance of the ED-based parametrization for both the protein and the nucleic acid. On the other hand, statistically significance differences concerning edProt were more difficult to interpret. edENM was not performing significantly better than edProt for *RMSIP* (*p*-value: ∼0.34), while it was close to a significant improvement for *O*_*max*_ (*p*-value: ∼0.08). On the other hand, edProt had a clear significant improvement over ENM in terms of *RMSIP* (*p*-value: ∼0.01) but not for *O*_*max*_ (*p*-value: ∼0.09). ENM was never found to perform better than either edProt or edENM (*p*-values > 0.9). Taken together, these results suggest that an ED-based parametrization both at the protein and nucleic acid level significantly improves the performance over the uniform-spring ENM, while the significancy at the level of protein and nucleic acid components separately might depend on a case-by-case basis. For example, edENM clearly outperformed both ENM and edProt in describing the PCs from the NMR ensemble of nucleolin RNA-binding domain 12 (RBD12) in complex with RNA (PDB: 1FJE). The experimental motion along PC1 was captured by edENM with 84% similarity with NM1, while the maximum agreement with ENM and edProt was found with NM3 with 66% and 60% similarity, respectively (Fig. 6). Looking at the predicted oscillations, we observed that the stronger nucleic acid backbone of edENM allows to avoid the generation of extreme displacements in the RNA tail that is observed in ENM and edProt (Fig. 6D). Together with the previous examples shown in Fig. 5 and Supplementary Fig. 11, this confirms that the ED-based parametrization improves the stability of the NMs in nucleic acids, avoiding backbone ruptures and extremely localized fluctuations.

**Figure 6.**
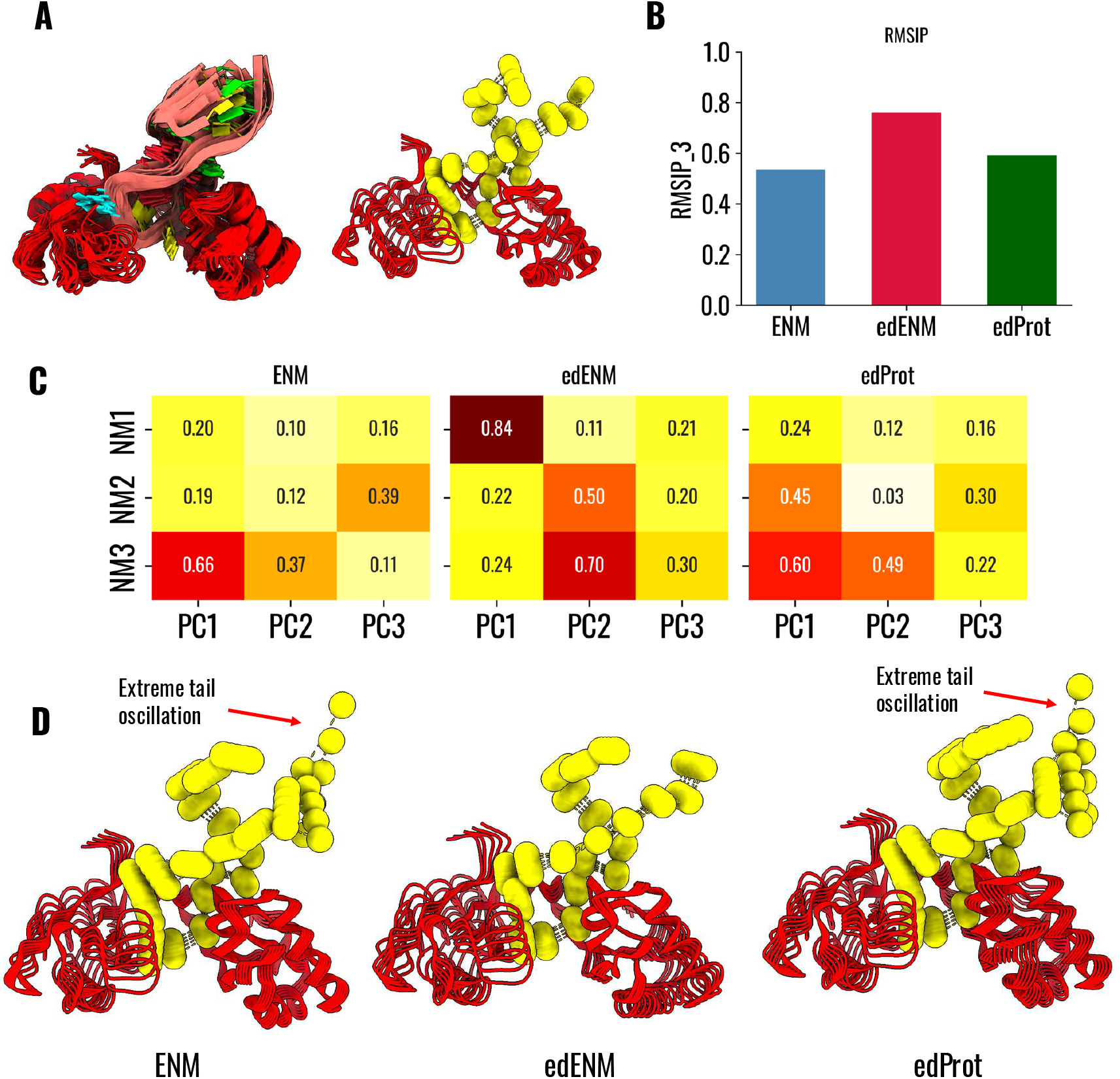
Comparison between ENM, edENM, and edProt and PC modes from the NMR ensemble (PDB: 1FJE) of the nucleolin RNA-binding domain 12 (RBD12) in complex with RNA. (A) Ensemble of 19 models (left) in the solution NMR used for PCA and direction of PC1 eigenvector (right), projected on the protein (red cartoon) and the RNA P-atoms (yellow spheres). (B) Comparison between ENM, edENM, and edProt in terms of *RMSIP* scores between the first 3 NMs and the first 3 experimental PCs. (C) Matrix of overlaps between the first 3 PC modes and the first 3 NMs from ENM, edENM, and edProt. (D) Graphical representation of the motions associated with NMs with the highest overlap with PC1, i.e. NM3 for ENM (left), NM1 for edENM (center), and NM3 for edProt. Note how all the models properly describe the conformational flexibility in the protein, but ENM and edProt introduce extreme oscillations in the RNA tail, which is not observed using the ED-refined nucleic-acid parametrization.

We also assessed the similarity of the NMs from edENM, ENM, and edProt to the experimental PC eigenvectors arising from a curated and highly heterogeneous set of protein-nucleic acid ensembles from X-ray and cryoEM conformations (Supplementary Table 4). In this case, we computed overlaps and *RMSIPs* considering the NMs from each individual PDB in the ensemble and Supplementary Table 10 reports average and maximum *O*_*max*_ and *RMSIP* scores for each ensemble. Overall, the similarity scores in edENM were found to surpass (∼ +1-4%) those from the ENM and edProt, confirming the slightly better performance of edENM in capturing the main directions of motions extracted from experimental ensembles. When assessing the statistical significance of such differences, these were found to be highly significant at the level of individual PDBs (*p*-values < 1 × 10^−4^, Supplementary Table 11) for both *O*_*max*_ and *RMSIP* (Supplementary Fig. 18). Due to the highly uneven number of PDB structures across different ensembles (Supplementary Table 4), we also assessed the statistical significance of these differences when considering aggregated *O*_*max*_ and *RMSIP* scores for each ensemble, aggregating by average and maximum values. In this case, the statistical significance of higher edENM scores was found to decrease both for average (Supplementary Table 12, Supplementary Fig. 19) and maximum scores (Supplementary Table 13, Supplementary Fig. 20). edENM was still found to perform significantly better than edProt (*p*-values < 0.05), suggesting the higher performance of employing an ED-based parametrization for the nucleic acid when coupled with an ED-based parametrization of the protein. ENM was never found to be significantly better than edENM (*p*-values ∼0.8) or edProt (*p*-values ∼0.4-0.9), while the opposite hypotheses (edENM > ENM and edProt > ENM) were associated with *p*-values of ∼0.2 for edENM and ∼0.05-0.6 for edProt.

For the entire dataset of PDBs of protein-nucleic acid ensembles, we also computed collectivity scores *κ* of the lowest-frequency normal mode (NM1) predicted by edENM, edProt, and ENM. NM1 represents the softest, lowest-energy mode and is generally expected to be highly collective. This analysis revealed that edENM consistently provides higher *κ* values (∼0.44 on average) than both edProt (∼0.36) and ENM (∼0.26), with highly significant differences in both cases (*p*-values < 1 × 10^−4^, Supplementary Table 14). As shown in Fig. 7, ENM can produce extremely localized vibrations, e.g., *κ* < 0.1, in the most dominant NM, while the ED-based parametrization generates consistently more collective motions, *κ* > 0.5. Taken together, these results confirm that edENM enhances the collectivity of the lowest-frequency harmonic vibrations.

**Figure 7.**
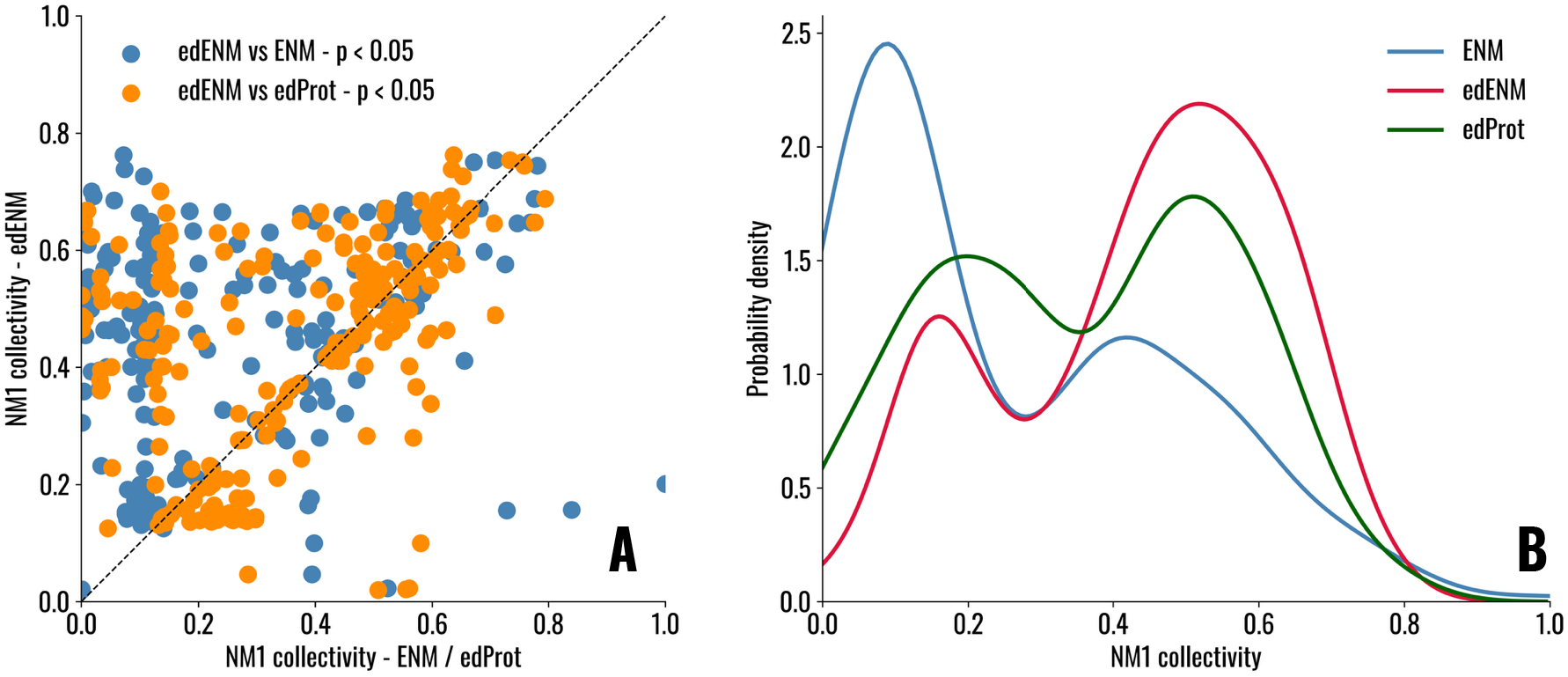
(A) Motion collectivity for the lowest-frequency normal mode (NM1) evaluated from edENM, ENM, and edProt. Scatterplot showing the comparison between NM1 collectivity between edENM and ENM (blue points) or edENM and edProt (orange points). (B) Distribution of NM1 collectivity values for ENM (blue), edENM (red), and edProt (green). *κ* values for NM1 averaged across the entire dataset are 0.436 ± 0.185 for edENM, 0.363 ± 0.201 for edProt, and 0.262 ± 0.222 for ENM. *P*-value analysis arising from one-sided Wilcoxon tests are reported in Supplementary Table 14.

### Exploring conformational transitions with eBDIMS

After assessing the performance of the edENM force-field for nucleic acids and protein-nucleic acid complexes via NMA, the optimal edENM parameters were integrated into the eBDIMS2 algorithm^49^, to be able to simulate transition pathways not only for proteins, but also for RNA, DNA, and any macromolecular complex.

A recent cryoEM study resolved the tertiary folds of the stem-loop 5 (SL5) genomic RNA elements in six coronaviruses, revealing a conserved T-shaped architecture composed of two perpendicular coaxial stacks formed by the stem-SL5c and SL5a-SL5b domains^59^. In two β-coronaviruses, *BtCoV-HKU5* and *MERS*, this architecture exhibits a pronounced conformational heterogeneity, characterized by a hinge-like motion originating from the SL5a internal loop^59^. Structural models of *BtCoV-HKU5* cluster into closed, open, and intermediate conformations (Fig. 8A). PCA of the experimental ensemble shows that the dominant motion – a large-scale hinge-bending transition between closed and open states – is captured by the first PC, accounting for ∼77% of the variance (Fig. 8B). NMA yielded very high overlaps between the lowest-frequency edENM modes and PC1 (*O*_*max*_ > 0.8), with edENM slightly outperforming the uniform-spring parametrization (Supplementary Table 6). To model the full hinge-bending transition (*RMSD* ∼10.5 Å) beyond the directionality of a small-scale harmonic vibration, we used eBDIMS to simulate both the closed-to-open and open-to-closed pathways. All simulations converged to the target states with final *RMSDs* below 1Å (Figs. 8B) and consistently approached the cluster of experimentally observed intermediate conformers before converging to the targets (Figs. 8C), demonstrating that eBDIMS can now generate conformational transitions not only in proteins^48,49^, but also in nucleic acids.

**Figure 8.**
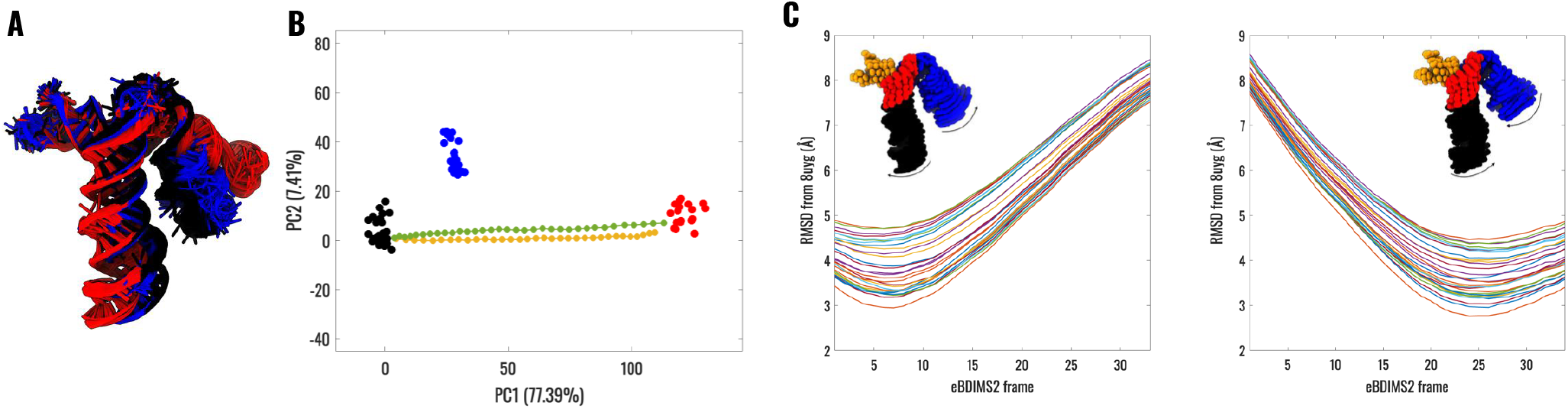
Conformational heterogeneity and eBDIMS transition pathways of the SL5 RNA from *BtCoV-HKU5* (∼40 kDa). (A) CryoEM SL5 conformations clustered into closed (black; PDB: 8UYE), open (red; 8UYJ), and intermediate (blue; 8UYG) states^59^. All deposited models within each PDB entry are shown to illustrate positional variability. (B) PCA projection of the experimental conformational ensemble and eBDIMS transition pathways (closed-to-open in orange; open-to-closed in green). (C) *RMSD* between eBDIMS frames and individual models of the intermediate conformer (8UYG). Each line refers to an individual model in the PDB file. Insets illustrate the hinge-bending motion of the SL5a arm along the eBDIMS trajectory, with P, C1’ and C2 atoms represented as spheres, and SL5 segments colored as follows: SL5-stem (black), SL5a (blue), SL5b (orange), SL5c (red).

Another recent cryoEM study identified two functional states of *A. Thaliana* Argonaute10 (AtAgo10) in complex with guide and target RNAs (Fig. 9): a slicing-incompetent bent-duplex state and a slicing-competent central-duplex state^60^. The transition between these states involves coordinated rearrangements of the PAZ and N domains^60^ and a localized reorganization of the bound RNA duplex. PCA of the two-state ensemble, including both protein and RNA components, yielded a single meaningful PC describing this transition, which spans a relevant structural change (*RMSD* ∼5.5 Å), but weakly collective (*κ* ∼ 0.07), as it mainly involves the repositioning of the RNA strands within the protein scaffold. Consistent with its localized and complex nature, this transition was poorly captured by NMA, with low overlaps between NMs and the transition vector (*O*_*max*_ ∼ 0.3), as low-frequency modes primarily describe collective motions of the whole protein-RNA complex (Fig. 9B-C). In contrast, eBDIMS could successfully be used to provide a mechanistic view of the complex repositioning of the two RNA strands from the bent-to the central-duplex state (Fig. 9D), reaching a final *RMSD* of ∼ 0.4 Å from the target structure. These results indicate that, although harmonic vibrational modes may fail to describe highly localized conformational changes^30,35^, ENM-based algorithms like eBDIMS can still allow to capture the underlying transition pathways, provide mechanistic insights, and generate conformational intermediates.

**Figure 9.**
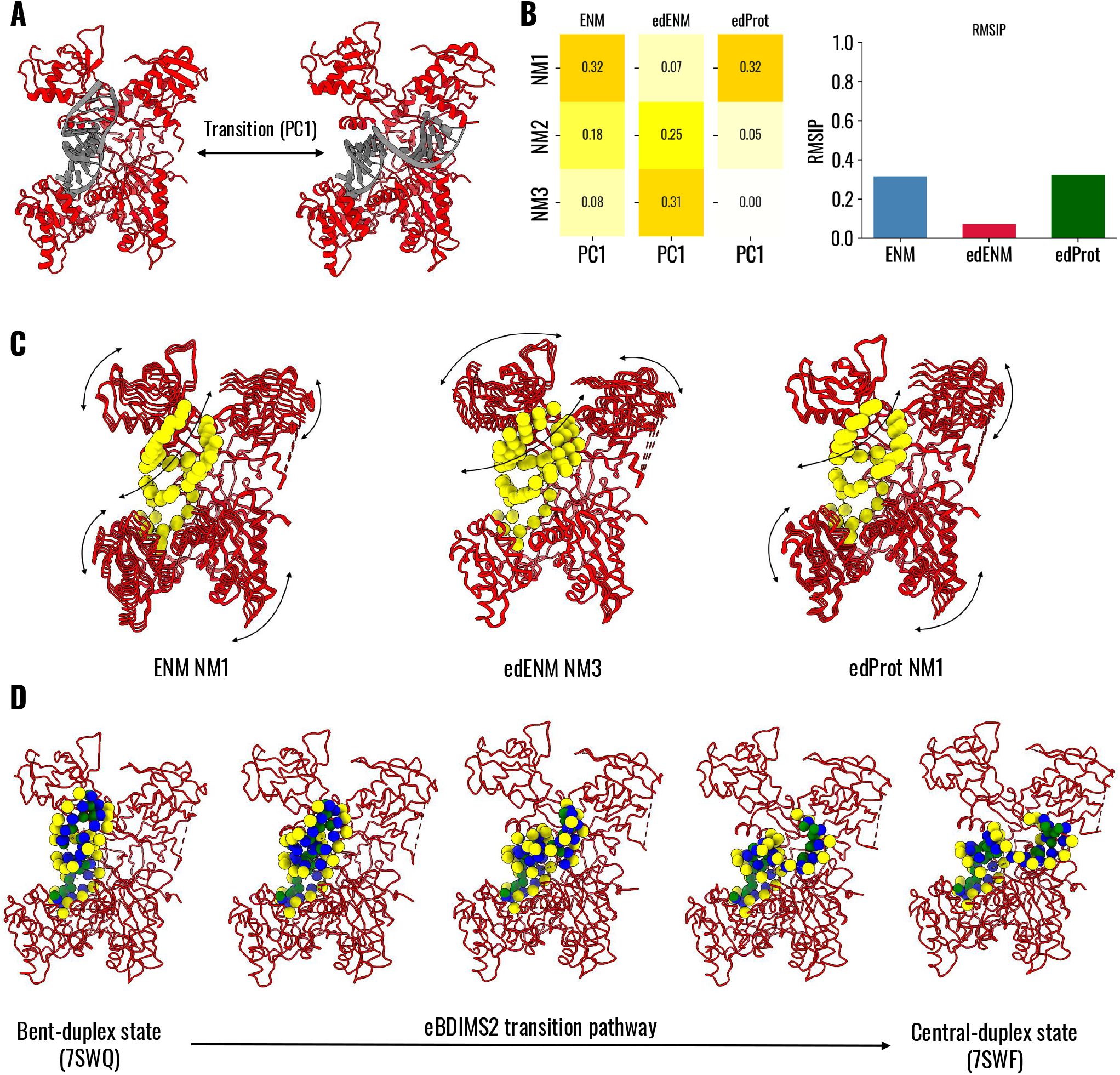
Conformational change and eBDIMS transition pathways of the *A. Thaliana* Argonaute10 (AtAgo10)-guide-target RNA complex (∼120 kDa). (A) CryoEM bent-duplex (PDB: 7SWQ, left) and central-duplex (7SWF, right) conformation of the AtAgo10–guide–target complex^60^, highlighting the motion of RNA chains within the complex (PC1). (B) Comparison between ENM, edENM, and edProt in terms of overlap and *RMSIP* scores between the first 3 NMs and conformational transition (PC1). (C) Graphical representation of the motions associated with NMs with the highest overlap with PC1, i.e. NM1 for ENM (left), NM3 for edENM (center), and NM1 for edProt. (D) Snapshots of AtAgo10– guide–target complex intermediates along the transition pathways simulated with eBDIMS from the bent-duplex to the central-duplex state, highlighting intermediate RNA positions.

As we recently demonstrated for protein systems, eBDIMS can be efficiently scaled to explore conformational transitions in macromolecular assemblies reaching the MDa scale^49^. Here, we further extend the applicability of the method to heterogeneous complexes composed of multiple proteins and long nucleic acid chains. The first example concerns a higher-order telomeric chromatin, which has been recently shown by cryoEM to adopt a columnar architecture characterized by tight nucleosome stacking^61^. Notably, these multi-nucleosome assemblies also populate alternative, more open conformational states, in which a single nucleosome disengages from the stack and flips outward^61^. Figure 10A illustrates structural intermediates generated by eBDIMS along the large-scale (*RMSD* ∼ 25 Å) closed-to-open transition of a telomeric trinucleosome. The resulting pathway revealed a highly cooperative rearrangement involving rigid-body translations and rotations of the moving histone core, coupled with DNA twisting motions, highlighting the complex mechanical coupling required to achieve the column unstacking. The second example addresses the nucleoplasmic maturation of a pre-40S ribosomal subunit, for which recent cryoEM studies have identified multiple assembly intermediates^62^. In an early nucloplasmic state (UTP14), the 3’ major domain of the 18S rRNA remains highly flexible and loosely connected to the remaining of the ribosomal body. In the subsequent RRP12-A1 state, the recruitment of additional assembly factors stabilizes the particle and repositions the 18S rRNA into a more stable complex^62^. eBDIMS simulations bridging these two end states revealed a pronounced hinge-bending motion of the 18S rRNA (Fig. 10B), in which most of the ribosomal subunit behaves quasi-rigidly while the rRNA pivots about a localized hinge region. This motion provides a mechanism for head-body reorganization during nuclear maturation, illustrating how localized RNA flexibility can enable global conformational changes also in larger assembly such as ribosomes.

**Figure 10.**
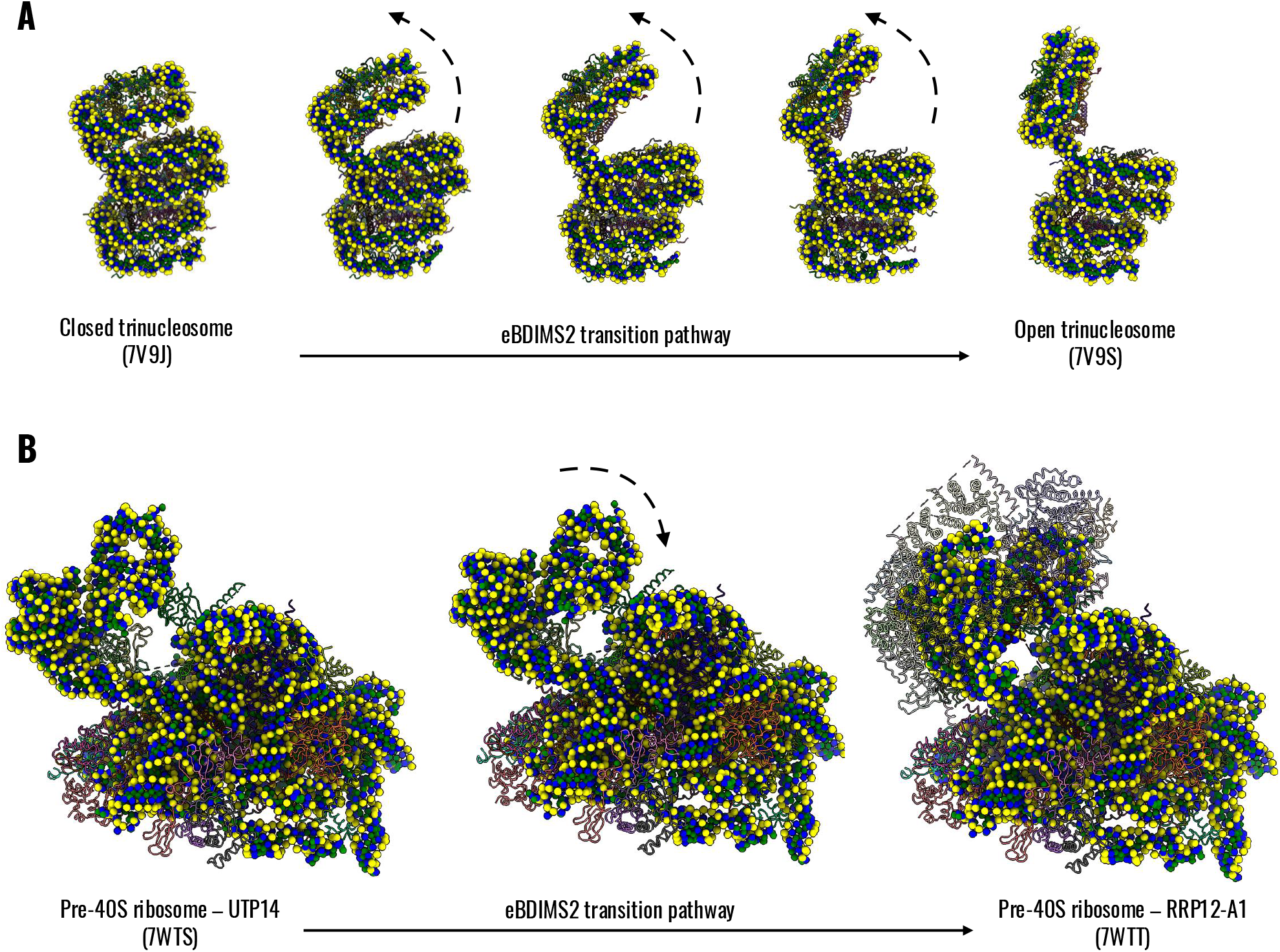
eBDIMS transition pathways of the (A) *H. Sapiens* telomeric trinucleosome (∼ 560 kDa) from closed columnar conformation (PDB: 7V9J) to open conformation (7V9S), highlighting the large-scale unstacking of one nucleosome; (B) *H. Sapiens* pre-40S ribosomal subunit (∼ 1.3 MDa) from UTP14 (7WTS) to RRP12-A1 state (7WTT), highlighting the hinge-bending motion of the 3’ domain of the large 18S rRNA and its stabilization into the larger assembly upon binding with the additional ribosomal proteins.

## Discussion

The development of ENMs for nucleic acids has so far been largely focused on RNA systems^37,39,40^, with comparatively limited attention devoted to DNA or unified descriptions. Early efforts explored the impact of CG strategies – such as the number and spatial placement of beads per nucleotide – on ENM performance, primarily evaluating their ability to reproduce atomic fluctuations or predict conformational changes^37^. More recently, ENMs for protein-nucleic acid complexes have been proposed in which parameter optimization was guided by maximizing the correlation between experimental and computed B-factors^41^. However, the use of B-factors as an optimization target remains controversial, as these are strongly influenced by crystal packing effects^44,45,63^. In contrast to the extensive methodological refinement achieved for protein ENMs^36^, the development of optimized and transferable ENMs for nucleic acids has remained comparatively underexplored.

In this work, we address this gap by introducing an MD-based parametrization of a nucleic acid ENM that employs a unified parameter set for both DNA and RNA, as well as for all types of protein-nucleic acid interactions. Our approach leverages extensive atomistic MD simulations together with experimentally observed conformational variability extracted from sets of PDB structures. Rather than targeting local fluctuations, the optimization protocol is explicitly designed to enhance the model’s ability to capture collective and large-scale conformational transitions observed experimentally. In doing so, we extend our previously developed edENM framework for proteins^31^ to nucleic acids and further to heterogeneous protein-nucleic acid assemblies. By benchmarking the method across a diverse set of experimental structures obtained by NMR, cryo-EM, and X-ray crystallography, and by direct comparison with a simpler (uniform-spring) ENM parametrization, we found that the proposed model achieves improved agreement with experimental motions (Fig. 4). Moreover, it yields more collective normal modes (Fig. 7), suppressing unphysical local backbone ruptures (Figs. 5-6) that arise from overly weak or poorly connected spring networks.

The novel integration of the optimized edENM into the eBDIMS framework^48,49^ enabled us not only to assess harmonic motions, but also to explore large-scale conformational transitions in a wide range of nucleic acid–containing systems, from isolated RNA folds to MDa-scale assemblies. eBDIMS could generate smooth transition pathways connecting experimentally resolved end states, providing structural intermediates and mechanistic insights even when the underlying motions were highly complex, such as the complex RNA rearrangement in AtAgo10 that was poorly described by harmonic NMA (Fig. 9). Thanks to our recent and more efficient implementation^49^, the approach could also be scaled to larger assemblies, as illustrated by the simulation of the telomeric trinucleosome and pre-40S ribosomal subunit (Fig. 10). Together, these results demonstrate that the proposed edENM can naturally extend the eBDIMS framework to nucleic acid and protein–nucleic acid systems, enabling the investigation of biologically relevant conformational transitions across a broad range of sizes and molecular architectures.

A major strength of the proposed framework lies in the intrinsic efficiency and interpretability of ENMs. ENMs provide a highly simplified yet physically grounded representation of macromolecular dynamics^64^, allowing conformational variability to be explored at a fraction of the cost of MD simulations, while still capturing dominant directions of motions^12,14^. Across the systems examined here, low-frequency NMs exhibited high overlaps with experimentally derived conformational changes, in some cases even higher than 90% (Fig. 5), underscoring the ability of topology-driven models to capture biologically relevant motions, without the need to embed additional chemistry information. It is well known that the performance of ENMs is strongly dictated by the network topology^33,34,36,65,66^, rather than the fine-tuning of force constants. This inherently limits the extent to which parameter optimization can yield dramatic gains in performance. Within this context, the improvements introduced by the present parametrization are nonetheless meaningful, as they address the main shortcomings of simpler uniform-spring models, including the suppression of unphysical backbone ruptures and overly localized modes (Figs. 5-7). By promoting more collective motions and enforcing a more coherent elastic coupling along the nucleic acid backbone, edENM provides a more robust description of large-scale, collective conformational changes.

Notably, the most pronounced performance gains were observed for nucleic acid–only systems, particularly NMR ensembles (Fig. 4), which are expected to reflect intrinsic dynamical fluctuations. This is consistent with the vibrational nature of ENMs and contrasts with protein–nucleic acid assemblies, where conformational differences from structural ensembles may not be only dictated by vibrational motions, but also by more complex, localized transitions – such as the RNA rearrangement observed in AtAgo10 (Fig. 9). Taken together, these observations highlight both the strengths and domains of applicability of ENM-based approaches.

Despite the performance of the proposed ENM framework, several general limitations should be acknowledged. First, the development and validation of the model was constrained by the limited availability of suitable structural datasets displaying large-scale conformational variability in nucleic acids and protein– nucleic acid systems. In total, our benchmarking relied on approximately ∼60 different macromolecular systems spanning both DNA, RNA, and protein-nucleic acid assemblies. While a higher number of available test cases would have strengthen the statistical significance of our validation, this data limitation reflects the broad imbalance in structural biology in which nucleic acids remain substantially underrepresented compared to proteins in public repositories such as the PDB^50^: currently, only ∼2% of PDB entries are nucleic acid-only, ∼6% comprise protein-nucleic acid systems, while ∼86% contain only protein entities. This scarcity is further highlighted by the difficulty of constructing coherent structural ensembles for complexes containing nucleic acids: even when complexes are captured in multiple conformational states, the associated nucleic acid components often differ in sequence or length, as in the case of RNaseIII-RNA complexes^48,67^, limiting their use in ensemble-based analyses.

A second important limitation concerns the structural resolution of CG models. While vibrational modes and eBDIMS-generated trajectories can provide valuable mechanistic insights into conformational changes at the CG (three-bead) level, approaches for reconstructing any CG model at atomistic resolution are still lacking for nucleic acids. Several reliable algorithms exist for CG–to–all-atom reconstruction of proteins, including *cg2all*^68^, *PULCHRA*^69^, etc., but tools for nucleic acids and protein–nucleic acid assemblies remain underdeveloped. Addressing this gap will be essential to re-introduce proper atomistic details to CG models containing nucleic acids for further downstream applications, such as MD simulations of conformational intermediates, interactions with small molecules, etc. This limitation also reflects a broader methodological imbalance in the field, where computational advances in structure prediction and conformational sampling have been overwhelmingly driven by protein-focused approaches, including deep-learning (DL) methods inspired by AlphaFold^70–74^. Some progress has been recently made for RNA^75–77^. However, the level of maturity and integration seen for protein systems has not been reached for nucleic acids yet. We believe that further methodological development, together with an increased experimental characterization of nucleic acid dynamics, is pivotal to fully unlock the potential of integrative modeling approaches for macromolecular assemblies including nucleic acids.

## Data Availability

All codes and data needed to reproduce the results shown in this work are freely available on GitHub at https://github.com/domenicoscaramozzino/Protein_DNA_RNA_edENM.

## Author Contributions

Conceptualization: M.C., D.S., M.A.D., L.O.; Methodology and model development: M.C., D.S., B.H.L.; Software implementation: M.C., D.S.; Data curation and computational analysis: M.C., D.S.; Validation and benchmarking: M.C., D.S.; Writing – original draft: M.C., D.S.; Writing – review and editing: M.C., D.S., B.H.L., M.A.D., L.O.; Supervision: M.A.D., L.O.; Funding acquisition: M.A.D., L.O.

## Acknowledgments

L.O. acknowledges financial support from Cancerfonden Junior Investigator Award (CF 21 0305 JIA) and Project Grants (CF 21 1471 Pj, CF 24 3801 Pj) as well as Vetenskapsrådet Starting Grant (VR 2021-02248) and Karolinska Institutet. D.S. acknowledges financial support from Cancerfonden Postdoctoral Fellowship (CF 24 0908 PT). M.A.D. acknowledges financial support from the European Union’s Horizon 2020 research and innovation program under the Marie Sklodowska-Curie Staff Exchange, through the GALATEA project (Grant Agreement No. 101183057).

## References

1. Bao, L., Zhang, X., Jin, L. & Tan, Z.-J. Flexibility of nucleic acids: From DNA to RNA. Chinese Phys. B 25, 018703 (2015).

2. Felsenfeld, G. & Groudine, M. Controlling the double helix. Nature 421, 448–453 (2003).

3. Sauerwald, N., Zhang, S., Kingsford, C. & Bahar, I. Chromosomal dynamics predicted by an elastic network model explains genome-wide accessibility and long-range couplings. Nucleic Acids Res 45, 3663–3673 (2017).

4. Peng, J., Yang, J., Anand, D. V., Shang, X. & Xia, K. Flexibility and rigidity index for chromosome packing, flexibility and dynamics analysis. Front. Comput. Sci. 16, 164902 (2021).

5. Vilarrasa-Blasi, R. et al. Dynamics of genome architecture and chromatin function during human B cell differentiation and neoplastic transformation. Nat Commun 12, 651 (2021).

6. Mustoe, A. M., Brooks, C. L. & Al-Hashimi, H. M. Hierarchy of RNA Functional Dynamics. Annu. Rev. Biochem. 83, 441–466 (2014).

7. Vicens, Q. & Kieft, J. S. Thoughts on how to think (and talk) about RNA structure. Proceedings of the National Academy of Sciences 119, e2112677119 (2022).

8. Assmann, S. M., Chou, H.-L. & Bevilacqua, P. C. Rock, scissors, paper: How RNA structure informs function. Plant Cell 35, 1671–1707 (2023).

9. Bose, R., Saleem, I. & Mustoe, A. M. Causes, functions, and therapeutic possibilities of RNA secondary structure ensembles and alternative states. Cell Chemical Biology 31, 17–35 (2024).

10. Gan, J. et al. Intermediate States of Ribonuclease III in Complex with Double-Stranded RNA. Structure 13, 1435–1442 (2005).

11. Rawat, N. & Biswas, P. Shape, flexibility and packing of proteins and nucleic acids in complexes. Phys. Chem. Chem. Phys. 13, 9632–9643 (2011).

12. Orellana, L. Large-Scale Conformational Changes and Protein Function: Breaking the in silico Barrier. Front. Mol. Biosci. 6, 117 (2019).

13. Velázquez-Muriel, J. A. et al. Comparison of molecular dynamics and superfamily spaces of protein domain deformation. BMC Struct Biol 9, 6 (2009).

14. Orellana, L. Are Protein Shape-Encoded Lowest-Frequency Motions a Key Phenotype Selected by Evolution? Applied Sciences 13, 6756 (2023).

15. Henzler-Wildman, K. A. et al. A hierarchy of timescales in protein dynamics is linked to enzyme catalysis. Nature 450, 913–916 (2007).

16. Marin-Gonzalez, A., Vilhena, J. G., Perez, R. & Moreno-Herrero, F. A molecular view of DNA flexibility. Quart. Rev. Biophys. 54, e8 (2021).

17. Hospital, A., Goñi, J. R., Orozco, M. & Gelpí, J. L. Molecular dynamics simulations: advances and applications. Adv Appl Bioinform Chem 8, 37–47 (2015).

18. Karplus, M. & Kuriyan, J. Molecular dynamics and protein function. Proc. Natl. Acad. Sci. U.S.A. 102, 6679–6685 (2005).

19. Šponer, J. et al. Molecular Dynamics Simulations of Nucleic Acids. From Tetranucleotides to the Ribosome. J. Phys. Chem. Lett. 5, 1771–1782 (2014).

20. Muscat, S., Martino, G., Manigrasso, J., Marcia, M. & De Vivo, M. On the Power and Challenges of Atomistic Molecular Dynamics to Investigate RNA Molecules. J. Chem. Theory Comput. acs.jctc.4c00773 (2024) doi:10.1021/acs.jctc.4c00773.

21. Daidone, I. & Amadei, A. Essential dynamics: foundation and applications. WIREs Comput Mol Sci 2, 762–770 (2012).

22. David, C. C. & Jacobs, D. J. Principal Component Analysis: A Method for Determining the Essential Dynamics of Proteins. Methods Mol Biol 1084, 193–226 (2014).

23. Sankar, K., Mishra, S. K. & Jernigan, R. L. Comparisons of Protein Dynamics from Experimental Structure Ensembles, Molecular Dynamics Ensembles, and Coarse-Grained Elastic Network Models. J. Phys. Chem. B 122, 5409–5417 (2018).

24. Mhashal, A. R., Yoluk, O. & Orellana, L. Exploring the Conformational Impact of Glycine Receptor TM1-2 Mutations Through Coarse-Grained Analysis and Atomistic Simulations. Front. Mol. Biosci. 9, 890851 (2022).

25. Fedulova, A. S. et al. Molecular dynamics simulations of nucleosomes are coming of age. WIREs Comput Mol Sci 14, e1728 (2024).

26. Bahar, I. & Rader, A. Coarse-grained normal mode analysis in structural biology. Current Opinion in Structural Biology 15, 586–592 (2005).

27. Tirion, M. M. Large amplitude elastic motions in proteins from a single-parameter, atomic analysis. Physical review letters 77, 1905 (1996).

28. Bahar, I., Atilgan, A. R. & Erman, B. Direct evaluation of thermal fluctuations in proteins using a single-parameter harmonic potential. Folding and Design 2, 173–181 (1997).

29. Atilgan, A. R. et al. Anisotropy of Fluctuation Dynamics of Proteins with an Elastic Network Model. Biophysical Journal 80, 505–515 (2001).

30. Tama, F. & Sanejouand, Y.-H. Conformational change of proteins arising from normal mode calculations. Protein Engineering, Design and Selection 14, 1–6 (2001).

31. Orellana, L. et al. Approaching Elastic Network Models to Molecular Dynamics Flexibility. J. Chem. Theory Comput. 6, 2910–2923 (2010).

32. Lopéz-Blanco, J. R., Garzón, J. I. & Chacón, P. iMod: multipurpose normal mode analysis in internal coordinates. Bioinformatics 27, 2843–2850 (2011).

33. Hoffmann, A. & Grudinin, S. NOLB: Nonlinear Rigid Block Normal-Mode Analysis Method. J. Chem. Theory Comput. 13, 2123–2134 (2017).

34. Khade, P. M. et al. hdANM: a new comprehensive dynamics model for protein hinges. Biophysical Journal 120, 4955–4965 (2021).

35. Scaramozzino, D., Piana, G., Lacidogna, G. & Carpinteri, A. Low-Frequency Harmonic Perturbations Drive Protein Conformational Changes. IJMS 22, 10501 (2021).

36. Park, S. W., Lee, B. H. & Kim, M. K. Elastic Network Model: A Coarse-Grained Approach to the Study of Biomolecular Dynamics. Multiscale Sci. Eng. 5, 104–118 (2023).

37. Setny, P. & Zacharias, M. Elastic Network Models of Nucleic Acids Flexibility. J. Chem. Theory Comput. 9, 5460–5470 (2013).

38. Afonin, K. A. et al. Computational and experimental characterization of RNA cubic nanoscaffolds. Methods 67, 256–265 (2014).

39. Zimmermann, M. T. & Jernigan, R. L. Elastic network models capture the motions apparent within ensembles of RNA structures. RNA 20, 792–804 (2014).

40. Pinamonti, G., Bottaro, S., Micheletti, C. & Bussi, G. Elastic network models for RNA: a comparative assessment with molecular dynamics and SHAPE experiments. Nucleic Acids Res 43, 7260–7269 (2015).

41. Procyk, J., Poppleton, E. & Šulc, P. Coarse-grained nucleic acid–protein model for hybrid nanotechnology. Soft Matter 17, 3586–3593 (2021).

42. Uyar, A., Kurkcuoglu, O., Nilsson, L. & Doruker, P. The elastic network model reveals a consistent picture on intrinsic functional dynamics of type II restriction endonucleases. Phys. Biol. 8, 056001 (2011).

43. Wang, Y. & Jernigan, R. L. Comparison of tRNA Motions in the Free and Ribosomal Bound Structures. Biophysical Journal 89, 3399–3409 (2005).

44. Carugo, O. & Argos, P. Reliability of atomic displacement parameters in protein crystal structures. Acta Crystallogr D Biol Crystallogr 55, 473–478 (1999).

45. Hinsen, K. Structural flexibility in proteins: impact of the crystal environment. Bioinformatics 24, 521–528 (2008).

46. Emperador, A., Carrillo, O., Rueda, M. & Orozco, M. Exploring the Suitability of Coarse-Grained Techniques for the Representation of Protein Dynamics. Biophysical Journal 95, 2127–2138 (2008).

47. Orellana, L., Gustavsson, J., Bergh, C., Yoluk, O. & Lindahl, E. eBDIMS server: protein transition pathways with ensemble analysis in 2D-motion spaces. Bioinformatics 35, 3505–3507 (2019).

48. Orellana, L., Yoluk, O., Carrillo, O., Orozco, M. & Lindahl, E. Prediction and validation of protein intermediate states from structurally rich ensembles and coarse-grained simulations. Nat Commun 7, 12575 (2016).

49. Scaramozzino, D., Lee, B. H. & Orellana, L. Efficient sampling of large-scale transition pathways and intermediate conformations in sub-mesoscopic protein complexes. Nat Commun 17, 2202 (2026).

50. Berman, H. M. et al. The Protein Data Bank. Nucleic Acids Research 28, 235–242 (2000).

51. Tiemann, J. K. S. et al. MDverse: Shedding Light on the Dark Matter of Molecular Dynamics Simulations. Preprint at 10.1101/2023.05.02.538537 (2023).

52. The UniProt Consortium. UniProt: the Universal Protein Knowledgebase in 2023. Nucleic Acids Research 51, D523–D531 (2023).

53. Zhang, C., Shine, M., Pyle, A. M. & Zhang, Y. US-align: universal structure alignments of proteins, nucleic acids, and macromolecular complexes. Nat Methods 19, 1109–1115 (2022).

54. Dykeman, E. C. & Sankey, O. F. Normal mode analysis and applications in biological physics. J Phys Condens Matter 22, 423202 (2010).

55. Scaramozzino, D., Lacidogna, G., Piana, G. & Carpinteri, A. A finite-element-based coarse-grained model for global protein vibration. Meccanica 54, 1927–1940 (2019).

56. Bochicchio, A. et al. Molecular basis for the increased affinity of an RNA recognition motif with re-engineered specificity: A molecular dynamics and enhanced sampling simulations study. PLOS Computational Biology 14, e1006642 (2018).

57. Sekulovski, S., Sušac, L., Stelzl, L. S., Tampé, R. & Trowitzsch, S. Structural basis of substrate recognition by human tRNA splicing endonuclease TSEN. Nat Struct Mol Biol 30, 834–840 (2023).

58. Kameda, T., Awazu, A. & Togashi, Y. Histone Tail Dynamics in Partially Disassembled Nucleosomes During Chromatin Remodeling. Front. Mol. Biosci. 6, (2019).

59. Kretsch, R. C. et al. Tertiary folds of the SL5 RNA from the 5′ proximal region of SARS-CoV-2 and related coronaviruses. Proc. Natl. Acad. Sci. U.S.A. 121, e2320493121 (2024).

60. Xiao, Y., Maeda, S., Otomo, T. & MacRae, I. J. Structural basis for RNA slicing by a plant Argonaute. Nat Struct Mol Biol 30, 778–784 (2023).

61. Soman, A. et al. Columnar structure of human telomeric chromatin. Nature 609, 1048–1055 (2022).

62. Cheng, J. et al. The nucleoplasmic phase of pre-40S formation prior to nuclear export. Nucleic Acids Res 50, 11924–11937 (2022).

63. Fuglebakk, E., Reuter, N. & Hinsen, K. Evaluation of Protein Elastic Network Models Based on an Analysis of Collective Motions. J Chem Theory Comput 9, 5618–5628 (2013).

64. Rader, A. Coarse-grained models: getting more with less. Current Opinion in Pharmacology 10, 753–759 (2010).

65. Giordani, G., Scaramozzino, D., Iturrioz, I., Lacidogna, G. & Carpinteri, A. Modal Analysis of the Lysozyme Protein Considering All-Atom and Coarse-Grained Finite Element Models. Applied Sciences 11, 547 (2021).

66. Koehl, P., Orland, H. & Delarue, M. Parameterizing elastic network models to capture the dynamics of proteins. Journal of Computational Chemistry 42, 1643–1661 (2021).

67. Gan, J. et al. Intermediate States of Ribonuclease III in Complex with Double-Stranded RNA. Structure 13, 1435–1442 (2005).

68. Heo, L. & Feig, M. One bead per residue can describe all-atom protein structures. Structure 32, 97-111.e6 (2024).

69. Rotkiewicz, P. & Skolnick, J. Fast procedure for reconstruction of full-atom protein models from reduced representations. J Comput Chem 29, 1460–1465 (2008).

70. del Alamo, D., Sala, D., Mchaourab, H. S. & Meiler, J. Sampling alternative conformational states of transporters and receptors with AlphaFold2. eLife 11, e75751 (2022).

71. Mirabello, C., Wallner, B., Nystedt, B., Azinas, S. & Carroni, M. Unmasking AlphaFold to integrate experiments and predictions in multimeric complexes. Nat Commun 15, 8724 (2024).

72. Kalakoti, Y. & Wallner, B. AFsample2 predicts multiple conformations and ensembles with AlphaFold2. Commun Biol 8, 373 (2025).

73. Cui, X. et al. Beyond static structures: protein dynamic conformations modeling in the post-AlphaFold era. Briefings in Bioinformatics 26, bbaf340 (2025).

74. Jumper, J. et al. Highly accurate protein structure prediction with AlphaFold. Nature 596, 583–589 (2021).

75. Wang, J., Fan, Y., Hong, L., Hu, Z. & Li, Y. Deep learning for RNA structure prediction. Current Opinion in Structural Biology 91, 102991 (2025).

76. Kagaya, Y. et al. NuFold: end-to-end approach for RNA tertiary structure prediction with flexible nucleobase center representation. Nat Commun 16, 881 (2025).

77. Patt, E., Classen, S., Hammel, M. & Schneidman-Duhovny, D. Predicting RNA structure and dynamics with deep learning and solution scattering. Biophysical Journal 124, 549–564 (2025).

