## Supplementary Material for "An essential dynamics-based elastic network model to unravel the conformational dynamics of DNA, RNA, and protein-nucleic acid complexes"

**Supplementary Table 1.** Details about the systems and MD simulations used in the study. The references to the publication and to the online data, number of replicas available, simulation length, simulating software, employed force-field, simulation temperature, and training-test split are indicated for each system.

| Simulation | Molecular system | # rep | Length [ns] | Software | Force-field | T [K] | Split |
|---|---|---|---|---|---|---|---|
| Vreede et al. Nucleic Acids Res. 2019[1,2] | DNA | 5 | 200 | GROMACS | AMBER03 | 300 | Train |
| Lemkul Nucleic Acids Res. 2020[3,4] | DNA | 3 | 1000 | OpenMM | CHARMM36 | 298 | Train |
| Lemkul Nucleic Acids Res. 2020[3,4] | DNA | 3 | 1000 | OpenMM | CHARMM36 | 298 | Train |
| Lemkul Nucleic Acids Res. 2020[3,4] | DNA | 3 | 1000 | OpenMM | Drude-2017 | 298 | Train |
| Lemkul Nucleic Acids Res. 2020[3,4] | DNA | 3 | 1000 | OpenMM | Drude-2017 | 298 | Test |
| Kameda et al. Phys. Rev. E 2021[5,6] | DNA | 10 | 50 | NAMD | CHARMM36 | 300 | Test |
| Pyne et al. Nat. Commun. 2021[7,8] | DNA | 8 | 30 | AMBER | AMBER99 | 300 | Test |
| Salsbury et al. ACS Omega 2022[4,9] | DNA | 3 | 1000 | OpenMM | CHARMM36 | 298 | Train |
| Salsbury et al. ACS Omega 2022[4,9] | DNA | 3 | 1000 | OpenMM | CHARMM36 | 298 | Train |
| Salsbury et al. ACS Omega 2022[4,9] | DNA | 3 | 2000 | OpenMM | CHARMM36 | 298 | Train |
| Salsbury et al. ACS Omega 2022[4,9] | DNA | 3 | 2000 | OpenMM | CHARMM36 | 298 | Train |
| Bochicchio et al. PLoS Comput. Biol. 2018[10,11] | RNA | 1 | 880 | GROMACS | AMBER19SB+OL3 | 298 | Train |
| Bochicchio et al. PLoS Comput. Biol. 2018[10,11] | RNA | 5 | 880 | AMBER | AMBER19SB+OL3 | 298 | Train |
| Bochicchio et al. PLoS Comput. Biol. 2018[10,12] | RNA | 3 | 1000 | AMBER | AMBER19SB+OL3 VdW mod | 298 | Train |
| Levintov and Vashisth Phys. Chem. Chem. Phys. 2021[13,14] | RNA | 14 | 2000 | AMBER | ff99OL3 | 300 | Train |

| | | | | | | | |
|---|---|---|---|---|---|---|---|
| De Bisschop et al. Non-coding RNA 2022[15,16] | RNA | 1 | 1000 | GROMACS | Amber ff99+ parmbsc0 | 300 | Test |
| Kameda et al. Front. Mol. Biosci. 2019[17,18] | Complex | 10 | 100 | NAMD | CHARMM36 | 310 | Test |
| Baltrukevich and Bartos Front. Chem. 2023[19,20] | Complex | 4 | 500 | AMBER | AMBER14SB+OL3 | 310 | Train |
| Baltrukevich and Bartos Front. Chem. 2023[19,20] | Complex | 4 | 500 | AMBER | AMBER19SB+OL3 | 310 | Train |
| Baltrukevich and Bartos Front. Chem. 2023[19,21] | Complex | 4 | 500 | AMBER | AMBER14SB+OL3 | 310 | Train |
| Baltrukevich and Bartos Front. Chem. 2023[19,21] | Complex | 4 | 500 | AMBER | AMBER19SB+OL3 | 310 | Train |
| Baltrukevich and Bartos Front. Chem. 2023[19,20] | Complex | 4 | 500 | DESMOND | OPLS4 | 300 | Train |
| Baltrukevich and Bartos Front. Chem. 2023[19,21] | Complex | 4 | 500 | DESMOND | OPLS4 | 300 | Train |
| Sekulovski et al. Nat. Struct. Mol. Biol. 2023[22,23] | Complex | 1 | 1000 | GROMACS | AMBER14+OL3 | 300 | Test |
| Riccardi et al. PLoS Comput. Biol. 2019[24,25] | Complex | 20 | 50 | GROMACS | AMBER03 | 300 | Train |
| Bochicchio et al. PLoS Comput. Biol. 2018[10,11] | Complex | 5 | 1000 | AMBER | AMBER19SB+OL3 | 298 | Test |

**Supplementary Table 2.** PDB IDs for the multi-model structures forming the experimental dataset for the nucleic acid edENM parametrization. The structures have been divided into training and test sets. Groups of PDB IDs forming a single ensemble are grouped into squared brackets.

| Split | PDB |
|---|---|
| Training set | 1A60, 1B36, 1JUA, 1P5O, 2LU0, 2N1Q, 7XH3, 8THV, [8UYK, 8UYL, 8UYM], [7UVT, 7XD3, 7XD4, 7XSK, 7XSL, 7XSM, 7YG8, 7YG9, 7YGA, 7YGB, 7YGC, 7YGD, 8HD6, 8HD7, 8I7N] |
| Testing set | 1AX6, 1BJ2, 1FYP, [1JOX, 1JP0], 2GV3, 2JTP, 2LI4, 2M12, 2M4Q, 2M8Z, 2MQT, 2MTJ, 2N6X, 2PCV, 5N5C, 5UZT, 6H1K, 8S1W, [8UYE, 8UYG, 8UYJ] |

**Supplementary Table 3.** PDB IDs for the NMR structures of protein-nucleic acid complexes used for final parameter refinement and performance testing.

| PDB |
|---|
| 1AHD, 1AUD, 1F6U, 1FJE, 1FNX, 1ULL, 2KEK, 2KM8, 2K1N, 2K7F, 2LT7, 2ME6, 2MF0, 2MKK, 2MRU, 2N3O, 2N82, 2OEH, 6GBM, 6WLH, 7WNR |

**Supplementary Table 4.** PDB IDs for structures in the experimental ensemble dataset for the testing of protein-nucleic complex edENM. The size of the molecular system is shown by reporting number of $C^{\alpha}$ atoms (for the protein) and P atoms (for DNA/RNA). Maximum *RMSD* values in each ensemble as well as the variances associated with PC1 and PC2 are also reported.

| Ensemble | Complex | PDB | Number $C^{\alpha}$/P atoms | RMSD (Å) | Variance PC1/PC2 (%) |
|---|---|---|---|---|---|
| 1 | HTH-type transcriptional regulator CueR bound to promoter DNA | 4WLS, 4WLW | 111/43 | 5.4 | 100.0/0.0 |
| 2 | DinG in complex with ssDNA | 6FWR, 6FWS | 683/9 | 4.4 | 100.0/0.0 |
| 3 | LigIV catalytic domain with bound DNA | 6BKF, 6BKG | 561/34 | 13.5 | 100.0/0.0 |
| 4 | Drosha and DGCR8 in complex with Primary MicroRNA | 6V5B, 6V5C | 1,301/66 | 11.4 | 100.0/0.0 |
| 5 | Ago10-guide-target RNA complex | 7SWF, 7SWQ | 753/31 | 5.5 | 100.0/0.0 |
| 6 | DNMT5 quaternary complex bound with DNA | 7R77, 7R78, 7T02 | 1,704/16 | 5.4 | 99.0/1.0 |
| 7 | dsRNA-binding protein Staufen homolog 1 bound with dsRNA | 6SDW, 6SDY (NMRs) | 70/33 | 4.8 | 58.3/9.0 |
| 8 | Flpb recombinase bound with DNA | 1FLO, 1M6X, 1P4E | 1,685/122 | 4.1 | 98.9/1.1 |
| 9 | Replicative DNA helicase bound with DNA | 7T20, 7T22, 7T23 | 2,661/10 | 5.6 | 75.7/24.3 |
| 10 | DNA topoisomerase 1 bound with DNA | 6CQ2, 6CQI, 8CZQ | 652/10 | 3.3 | 96.8/3.2 |
| 11 | Group II-encoded protein LTRA bound with Group II-intron RNA | 5G2X, 7D1A, 8H2H | 426/700 | 4.7 | 89.9/10.1 |
| 12 | 50S ribosomal protein L7Ae bound with RNA | 5DCV_1, 5DCV_2, 5XTM_1, 5XTM_2, 5Y7M_1, 5Y7M_2 | 118/46 | 9.4 | 67.4/29.9 |

| | | | | | |
|---|---|---|---|---|---|
| 13 | Gag-Pro-Pol polyprotein bound with DNA | 5EJK, 7JN3, 7KU7, 7KUI, 8E14 | 1,158/64 | 3.3 | 60.5/35.3 |
| 14 | DNA primase/helicase bound with DNA | 6N7N, 6N7S, 6N7T | 1,597/13 | 7.4 | 93.0/7.0 |
| 15 | CRISPR-associated endonuclease C2c1 bound with DNA and RNA | 5U30, 5U31, 5U33, 5U34, 5WQE | 932/59 | 3.5 | 91.6/8.2 |
| 16 | Rad2p bound with DNA | 4Q0Z_1, 4Q0Z_2, 4Q10 | 588/30 | 3.9 | 92.1/7.9 |
| 17 | DNA polymerase delta bound with DNA | 6S1M, 6S1N, 6TNY, 6TNZ | 2,399/49 | 5.5 | 95.9/4.1 |
| 18 | CRISPR-associated endonuclease Cas9 bound with RNA | 6JDQ, 6JE9, 6KC7, 6KC8, 8HJ4, 8JA0 | 896/110 | 16.1 | 86.2/10.2 |
| 19 | DNA polymerase gamma 1/2 bound with DNA | 4ZTU, 4ZTZ, 5C51, 5C52, 5C53, 8D33, 8D37, 8D3R, 8D42 | 1,643/41 | 7.3 | 68.5/24.1 |
| 20 | DNA polymerase beta bound with DNA | 1HUO, 1HUZ, 2BPF, 2BPG | 322/13 | 6.6 | 67.8/30.3 |
| 21 | DNA polymerase beta bound with DNA | 3UXP, 3V72, 3V7J, 3V7K | 326/14 | 4.0 | 92.7/3.8 |
| 22 | Lactose operon repressor bound with DNA | 2KEI, 2KEJ, 2KEK (NMRs) | 104/45 | 5.4 | 53.1/19.1 |
| 23 | Human topoisomerase IIbeta in complex with DNA | 3QX3, 4G0U, 4G0V, 4G0W, 4J3N, 5GWI, 5GWJ, 5ZAD | 1,308/39 | 5.0 | 88.1/8.3 |
| 24 | Replication factor C bound with DNA | 7TFH, 7TFI, 7TFJ, 7TIB, 7TID | 2,446/38 | 11.5 | 98.3/0.9 |
| 25 | crRNA-guided Csy surveillance complex | 6B45, 6B47, 6B48, 6NE0, 6VQV, 7ECV, 7JZW, 7JZX, 7JZY, 7JZZ | 2,487/57 | 12.1 | 83.3/6.0 |
| 26 | CRISPR-associated endonuclease Cas9/Csn1 bound with DNA and RNA | 4OO8, 4UN3, 4UN4, 4UN5, 4ZT0, 4ZT9, 5B2R, 5B2S, 5B2T, 5FQ5, 5FW1, 5FW2, 5FW3, 5VW1, 5XBL, 6AEG, 6AI6, 6K3Z, 6K4P, 6K4Q, 6K4S, 6K4U, 6K57, 6O0Z, 6VPC, 7OX7, 7OX8, 7OX9, 7OXA, 7QQO, 7QQP, 7QQQ, 7QQR, 7QQS, 7QQT, 7QQU, 7QQV, 7QQW, 7QQX, 7QQZ, 7QR0, 7QR1, 7QR5, 7QR7, 7QR8, 7S36, 7S38, 7S4X, 7Z4C, 7Z4E, 7Z4H, 7Z4I, 7Z4J, 7Z4K, 7Z4L, 8G1I | 1,110/61 | 14.4 | 56.6/18.9 |

**Supplementary Table 5.** Set of parameters employed in the random search optimization for the nucleic acids edENM. Numbers in **bold** represent those selected as optimal after refinement.

| **Bond type** | **Covalent (pyrimidines)** | **Covalent (purines)** | **Hbond** | **Pseudo-covalent** | **Van der Waals** |
|---|---|---|---|---|---|
| Distance-dependent decay | No | No | No | Yes | Yes |
| *n* | - | - | - | 2.7, **2.8**, 2.9, 3.0 | **1.0**, 1.1, 1.2, 1.3 |
| *C* | 250, 270, **290**, 310, 330, 350 | 100, **120**, 140, 160, 180, 200 | 60, **65**, 70, 75, 80, 85 | 12, 14, 16, 18, **20** | **25**, 30, 35, 40 |
| *Cutoff* | - | - | - | 9, 10, **11**, 12, 13, 14 | 9, 10, **11**, 12, 13, 14 |

**Supplementary Table 6.** Testing of edENM and comparison against uniform-spring ENM for the testing set of nucleic acid multi-model PDB ensembles in Supplementary Table 2 in terms of overlaps and *RMSIP* scores between NMs and experimental PCs. Maximum overlaps $O_{max}$ and *RMSIP* were computed based on the first 3 NMs and 3 PCs. The numbers of the NM and PC vector giving the maximum overlap are indicated in the first and second position of the parenthesis, respectively. The last rows show average and standard deviations for all tested PDB structures as well as *p*-values assessing the statistical significance of the differences.

| **PDB** | $O_{max}^{ENM}$ | $O_{max}^{edENM}$ | $RMSIP^{ENM}$ | $RMSIP^{edENM}$ |
|---|---|---|---|---|
| 1AX6 | 0.583^(3,1) | 0.738^(1,1) | 0.528 | 0.553 |
| 1BJ2 | 0.713^(1,2) | 0.621^(1,2) | 0.690 | 0.653 |
| 1FYP | 0.769^(1,1) | 0.744^(2,2) | 0.772 | 0.893 |
| 1JOX | 0.728^(1,2) | 0.720^(3,1) | 0.704 | 0.634 |
| 1JP0 | 0.913^(1,1) | 0.736^(1,1) | 0.657 | 0.670 |
| 2GV3 | 0.576^(2,1) | 0.600^(1,3) | 0.532 | 0.564 |
| 2JTP | 0.566^(2,1) | 0.658^(1,1) | 0.557 | 0.679 |
| 2LI4 | 0.852^(2,2) | 0.892^(2,2) | 0.911 | 0.914 |
| 2M12 | 0.745^(3,2) | 0.607^(2,2) | 0.604 | 0.618 |
| 2M4Q | 0.783^(2,1) | 0.819^(1,1) | 0.707 | 0.721 |
| 2M8Z | 0.797^(1,1) | 0.769^(1,1) | 0.810 | 0.806 |
| 2MQT | 0.628^(2,1) | 0.797^(2,1) | 0.530 | 0.695 |
| 2MTJ | 0.852^(2,1) | 0.737^(1,1) | 0.594 | 0.675 |
| 2N6X | 0.904^(1,1) | 0.907^(1,1) | 0.793 | 0.808 |
| 2PCV | 0.636^(1,1) | 0.930^(1,1) | 0.570 | 0.758 |
| 5N5C | 0.950^(1,1) | 0.926^(1,1) | 0.609 | 0.628 |
| 5UZT | 0.603^(1,1) | 0.659^(2,1) | 0.613 | 0.621 |
| 6H1K | 0.779^(1,2) | 0.723^(1,2) | 0.616 | 0.687 |
| 8S1W | 0.775^(1,1) | 0.726^(1,1) | 0.707 | 0.712 |
| 8UYE | 0.785^(3,1) | 0.805^(2,1) | 0.515 | 0.661 |
| 8UYG | 0.798^(1,1) | 0.858^(1,1) | 0.632 | 0.666 |
| 8UYJ | 0.804^(1,1) | 0.858^(1,1) | 0.581 | 0.691 |
| All | 0.752 ± 0.110 | 0.765 ± 0.099 | 0.647 ± 0.102 | 0.696 ± 0.090 |
| Wilcoxon one-sided *p*-value (edENM > ENM) | 0.3278 | | 0.0004 | |

**Supplementary Table 7.** Set of parameters for the random search optimization of protein-nucleic acid interactions. Numbers in **bold** represent those selected as optimal after refinement.

| Bond type | Strong | Weak |
|---|---|---|
| Distance-dependent decay | Yes | Yes |
| *n* | 1.9, 2.0, 2.1, **2.2** | **1.0** |
| *C* | 15, 20, **25** | **30**, 35, 40 |
| *Cutoff* | **8**, 9 | **8**, 9 |

**Supplementary Table 8.** Testing of edENM and comparison against uniform-spring ENM and edProt for the set of protein-nucleic acid NMR models in Supplementary Table 3 in terms of overlap and *RMSIP* scores between NMs and experimental PCs. Maximum overlaps $O_{max}$ and *RMSIP* were computed based on the first 3 NMs and 3 PCs. The numbers of the NM and PC vector giving the maximum overlap are indicated in the first and second position of the parenthesis, respectively. The last row shows average and standard deviations for all tested PDB structures.

| PDB | $O_{max}^{ENM}$ | $O_{max}^{edProt}$ | $O_{max}^{edENM}$ | $RMSIP_3^{ENM}$ | $RMSIP_3^{edProt}$ | $RMSIP_3^{edENM}$ |
|---|---|---|---|---|---|---|
| 1AHD | $0.551^{(2,2)}$ | $0.457^{(2,2)}$ | $0.408^{(2,1)}$ | 0.415 | 0.425 | 0.393 |
| 1AUD | $0.717^{(1,2)}$ | $0.714^{(1,2)}$ | $0.622^{(1,1)}$ | 0.674 | 0.659 | 0.555 |
| 1F6U | $0.419^{(2,2)}$ | $0.492^{(3,3)}$ | $0.628^{(2,3)}$ | 0.300 | 0.561 | 0.567 |
| 1FJE | $0.656^{(3,1)}$ | $0.603^{(3,1)}$ | $0.842^{(1,1)}$ | 0.535 | 0.591 | 0.760 |
| 1FNX | $0.332^{(1,2)}$ | $0.682^{(3,1)}$ | $0.752^{(3,1)}$ | 0.282 | 0.514 | 0.480 |
| 1ULL | $0.544^{(1,1)}$ | $0.542^{(1,1)}$ | $0.620^{(1,1)}$ | 0.536 | 0.538 | 0.420 |
| 2KEK | $0.555^{(3,1)}$ | $0.549^{(3,3)}$ | $0.607^{(1,1)}$ | 0.569 | 0.582 | 0.599 |
| 2KM8 | $0.739^{(1,1)}$ | $0.876^{(1,1)}$ | $0.665^{(1,1)}$ | 0.535 | 0.586 | 0.550 |
| 2K1N | $0.251^{(2,2)}$ | $0.369^{(3,2)}$ | $0.447^{(2,2)}$ | 0.204 | 0.320 | 0.366 |
| 2K7F | $0.423^{(2,1)}$ | $0.452^{(2,1)}$ | $0.619^{(1,2)}$ | 0.320 | 0.337 | 0.522 |
| 2LT7 | $0.778^{(2,1)}$ | $0.791^{(1,1)}$ | $0.664^{(1,1)}$ | 0.496 | 0.511 | 0.516 |
| 2ME6 | $0.211^{(3,2)}$ | $0.292^{(3,2)}$ | $0.343^{(1,1)}$ | 0.194 | 0.269 | 0.278 |
| 2MF0 | $0.807^{(2,1)}$ | $0.794^{(1,1)}$ | $0.752^{(1,1)}$ | 0.552 | 0.559 | 0.563 |
| 2MKK | $0.503^{(2,1)}$ | $0.459^{(3,1)}$ | $0.689^{(1,1)}$ | 0.579 | 0.536 | 0.570 |
| 2MRU | $0.378^{(2,1)}$ | $0.345^{(2,1)}$ | $0.513^{(2,1)}$ | 0.309 | 0.346 | 0.434 |
| 2N30 | $0.560^{(1,2)}$ | $0.561^{(1,2)}$ | $0.790^{(2,1)}$ | 0.608 | 0.610 | 0.685 |
| 2N82 | $0.492^{(3,3)}$ | $0.515^{(1,1)}$ | $0.359^{(1,1)}$ | 0.530 | 0.425 | 0.370 |
| 2OEH | $0.216^{(3,2)}$ | $0.339^{(3,3)}$ | $0.322^{(2,3)}$ | 0.217 | 0.344 | 0.359 |
| 6GBM | $0.659^{(1,2)}$ | $0.722^{(2,1)}$ | $0.767^{(1,1)}$ | 0.629 | 0.723 | 0.648 |
| 6WLH | $0.632^{(2,2)}$ | $0.562^{(2,1)}$ | $0.486^{(1,2)}$ | 0.626 | 0.533 | 0.486 |
| 7WNR | $0.385^{(2,2)}$ | $0.406^{(2,2)}$ | $0.482^{(3,2)}$ | 0.385 | 0.406 | 0.482 |
| All | 0.515 ± 0.175 | 0.550 ± 0.159 | 0.590 ± 0.151 | 0.452 ± 0.152 | 0.494 ± 0.120 | 0.505 ± 0.115 |

**Supplementary Table 9.** *P*-value statistical test (Wilcoxon one-sided) between edENM, ENM, and edProt for $O_{max}$ values (left) and *RMSIP* scores (right) obtained from the dataset of protein-nucleic acid NMR structures from Supplementary Table 8. The matrix shows one-sided *p*-values with the hypothesis that the model in the column provides values significantly higher than the model in the row.

| Model | ENM | edProt | edENM | ENM | edProt | edENM |
|---|---|---|---|---|---|---|
| ENM | - | 0.0895 | 0.0210 | - | 0.0097 | 0.0273 |
| edProt | 0.9161 | - | 0.0786 | 0.9912 | - | 0.3414 |
| edENM | 0.9808 | 0.9265 | - | 0.9749 | 0.6711 | - |

**Supplementary Table 10.** Testing of edENM and comparison against ENM and edProt for the testing set of protein-nucleic acid ensembles in Supplementary Table 3 in terms of overlaps and *RMSIP* scores between NMs and experimental PCs. $O_{max}$ and *RMSIP* scores were computed based on the first 3 NMs and the maximum between 3 and the number of available experimental PCs. Each row reports average (left) and maximum (right) scores considering individual computations for each PDB in the ensemble. The last two rows report average scores and standard deviations for all tested ensembles considering aggregated scores by average or maximum values.

| **Ensemble** | $O_{max}^{ENM}$ | $O_{max}^{edProt}$ | $O_{max}^{edENM}$ | $RMSIP^{ENM}$ | $RMSIP^{edProt}$ | $RMSIP^{edENM}$ |
|---|---|---|---|---|---|---|
| 1 | 0.366/0.367 | 0.390/0.491 | 0.634/0.702 | 0.271/0.367 | 0.076/0.143 | 0.191/0.203 |
| 2 | 0.634/0.684 | 0.635/0.684 | 0.622/0.669 | 0.634/0.684 | 0.635/0.684 | 0.622/0.669 |
| 3 | 0.406/0.500 | 0.327/0.390 | 0.396/0.502 | 0.105/0.165 | 0.227/0.390 | 0.323/0.502 |
| 4 | 0.582/0.604 | 0.577/0.613 | 0.568/0.603 | 0.141/0.195 | 0.287/0.541 | 0.282/0.533 |
| 5 | 0.278/0.317 | 0.254/0.323 | 0.244/0.312 | 0.238/0.317 | 0.254/0.323 | 0.125/0.177 |
| 6 | 0.340/0.401 | 0.376/0.453 | 0.375/0.439 | 0.213/0.303 | 0.263/0.297 | 0.280/0.323 |
| 7 | 0.619/0.705 | 0.640/0.740 | 0.579/0.825 | 0.581/0.670 | 0.621/0.728 | 0.506/0.638 |
| 8 | 0.680/0.730 | 0.539/0.554 | 0.552/0.566 | 0.570/0.583 | 0.497/0.512 | 0.483/0.494 |
| 9 | 0.459/0.485 | 0.448/0.456 | 0.444/0.455 | 0.462/0.497 | 0.466/0.500 | 0.465/0.499 |
| 10 | 0.597/0.600 | 0.619/0.625 | 0.619/0.629 | 0.441/0.445 | 0.483/0.488 | 0.489/0.493 |
| 11 | 0.098/0.120 | 0.081/0.094 | 0.088/0.095 | 0.074/0.086 | 0.065/0.075 | 0.065/0.085 |
| 12 | 0.502/0.584 | 0.515/0.585 | 0.565/0.621 | 0.462/0.562 | 0.470/0.562 | 0.476/0.576 |
| 13 | 0.421/0.427 | 0.445/0.468 | 0.437/0.490 | 0.445/0.471 | 0.510/0.523 | 0.410/0.538 |
| 14 | 0.447/0.532 | 0.379/0.422 | 0.368/0.426 | 0.447/0.481 | 0.401/0.445 | 0.395/0.429 |
| 15 | 0.558/0.680 | 0.574/0.736 | 0.571/0.730 | 0.409/0.477 | 0.433/0.495 | 0.437/0.499 |
| 16 | 0.900/0.967 | 0.764/0.781 | 0.877/0.908 | 0.765/0.779 | 0.753/0.789 | 0.783/0.788 |
| 17 | 0.888/0.937 | 0.616/0.650 | 0.772/0.875 | 0.595/0.615 | 0.582/0.587 | 0.575/0.583 |
| 18 | 0.368/0.720 | 0.592/0.780 | 0.682/0.812 | 0.271/0.498 | 0.494/0.677 | 0.637/0.739 |
| 19 | 0.286/0.659 | 0.503/0.564 | 0.553/0.605 | 0.254/0.506 | 0.445/0.505 | 0.534/0.547 |
| 20 | 0.733/0.814 | 0.830/0.877 | 0.837/0.891 | 0.618/0.644 | 0.637/0.643 | 0.634/0.642 |
| 21 | 0.504/0.525 | 0.527/0.566 | 0.546/0.575 | 0.461/0.553 | 0.505/0.562 | 0.605/0.641 |
| 22 | 0.664/0.792 | 0.704/0.795 | 0.722/0.815 | 0.697/0.783 | 0.674/0.742 | 0.757/0.794 |
| 23 | 0.473/0.535 | 0.449/467 | 0.425/0.621 | 0.424/0.469 | 0.400/0.420 | 0.391/0.527 |
| 24 | 0.400/0.576 | 0.421/0.767 | 0.387/0.580 | 0.325/0.406 | 0.336/0.500 | 0.321/0.527 |
| 25 | 0.315/0.468 | 0.277/0.358 | 0.312/0.386 | 0.265/0.436 | 0.258/0.297 | 0.277/0.305 |
| 26 | 0.326/0.612 | 0.451/0.657 | 0.486/0.678 | 0.303/0.584 | 0.424/0.544 | 0.454/0.565 |
| All (avg.) | 0.494 ± 0.185 | 0.497 ± 0.162 | 0.525 ± 0.177 | 0.403 ± 0.181 | 0.431 ± 0.170 | 0.443 ± 0.176 |
| All (max.) | 0.590 ± 0.184 | 0.573 ± 0.176 | 0.608 ± 0.188 | 0.484 ± 0.170 | 0.499 ± 0.169 | 0.512 ± 0.172 |

**Supplementary Table 11.** *P*-value statistical test (Wilcoxon one-sided) between edENM, ENM, and edProt for $O_{max}$ values (left) and *RMSIP* scores (right) obtained from the dataset of protein-nucleic acid ensembles, whose aggregated values are reported in Supplementary Table 10. These *p*-value results have been obtained considering comparisons between similarity scores at the level of each individual PDB. The matrix shows one-sided *p*-values with the hypothesis that the model in the column provides values significantly higher than the model in the row.

| **Model** | ENM | edProt | edENM | ENM | edProt | edENM |
|---|---|---|---|---|---|---|
| ENM | - | $< 1 \times 10^{-6}$ | $< 1 \times 10^{-6}$ | - | $< 1 \times 10^{-6}$ | $< 1 \times 10^{-6}$ |
| edProt | 1.000 | - | 0.00008 | 1.000 | - | 0.0001 |
| edENM | 1.000 | 0.9999 | - | 1.000 | 0.9999 | - |

**Supplementary Table 12.** *P*-value statistical test (Wilcoxon one-sided) between edENM, ENM, and edProt for $O_{max}$ values (left) and *RMSIP* scores (right) obtained from the dataset of protein-nucleic acid ensembles, whose aggregated values are reported in Supplementary Table 10. These *p*-value results have been obtained considering comparisons between similarity scores aggregated through average values at the level of each ensemble. The matrix shows one-sided *p*-values with the hypothesis that the model in the column provides values significantly higher than the model in the row.

| **Model** | ENM | edProt | edENM | ENM | edProt | edENM |
|---|---|---|---|---|---|---|
| ENM | - | 0.34 | 0.23 | - | 0.05 | 0.12 |
| edProt | 0.66 | - | 0.04 | 0.95 | - | 0.20 |
| edENM | 0.77 | 0.96 | - | 0.88 | 0.80 | - |

**Supplementary Table 13.** *P*-value statistical test (Wilcoxon one-sided) between edENM, ENM, and edProt for $O_{max}$ values (left) and *RMSIP* scores (right) obtained from the dataset of protein-nucleic acid ensembles, whose aggregated values are reported in Supplementary Table 10. These *p*-value results have been obtained considering comparisons between similarity scores aggregated through maximum values at the level of each ensemble. The matrix shows one-sided *p*-values with the hypothesis that the model in the column provides values significantly higher than the model in the row.

| **Model** | ENM | edProt | edENM | ENM | edProt | edENM |
|---|---|---|---|---|---|---|
| ENM | - | 0.56 | 0.18 | - | 0.35 | 0.19 |
| edProt | 0.42 | - | 0.003 | 0.65 | - | 0.04 |
| edENM | 0.82 | 0.997 | - | 0.81 | 0.96 | - |

**Supplementary Table 14.** *P*-value statistical test (Wilcoxon one-sided) between edENM, ENM, and edProt for NM1 collectivity values obtained from the dataset of protein-nucleic acid ensembles, whose aggregated values are reported in Supplementary Table 10. These *p*-value results have been obtained considering comparisons between similarity scores at the level of each individual PDB. The matrix shows one-sided *p*-values with the hypothesis that the model in the column provides values significantly higher than the model in the row.

| **Model** | ENM | edProt | edENM |
|---|---|---|---|
| ENM | - | $< 1 \times 10^{-6}$ | $< 1 \times 10^{-6}$ |
| edProt | 1.000 | - | 0.00007 |
| edENM | 1.000 | 0.9999 | - |

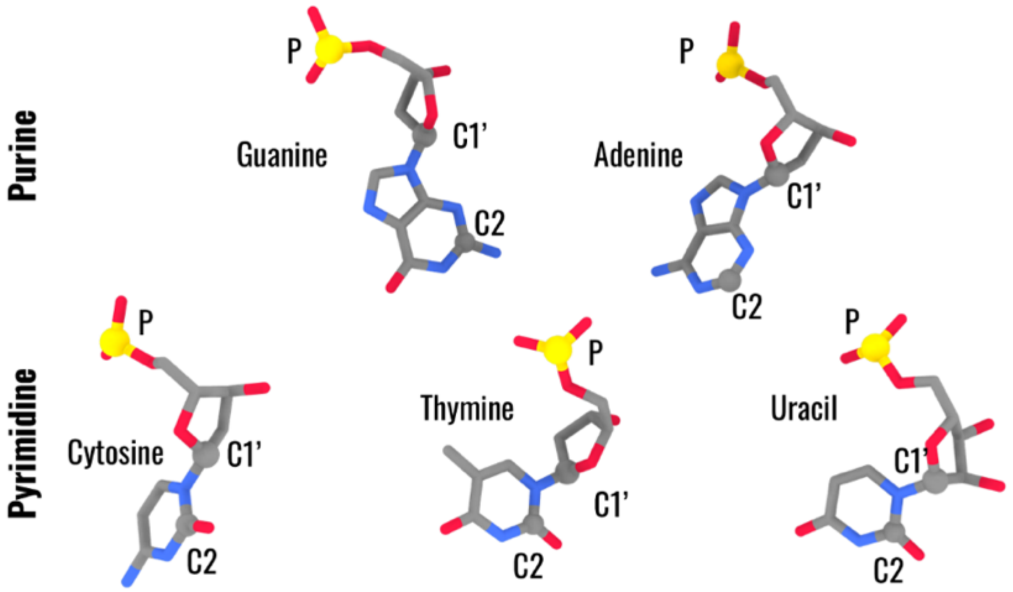


**Supplementary Figure 1.** Representation of the three-bead SBP model from Pinamonti et al.[26] for the five nucleotide types (G, A, C, T, U). The three beads of the ENM, shown as spheres, are located in the geometrical positions of P, C1', and C2 atoms.

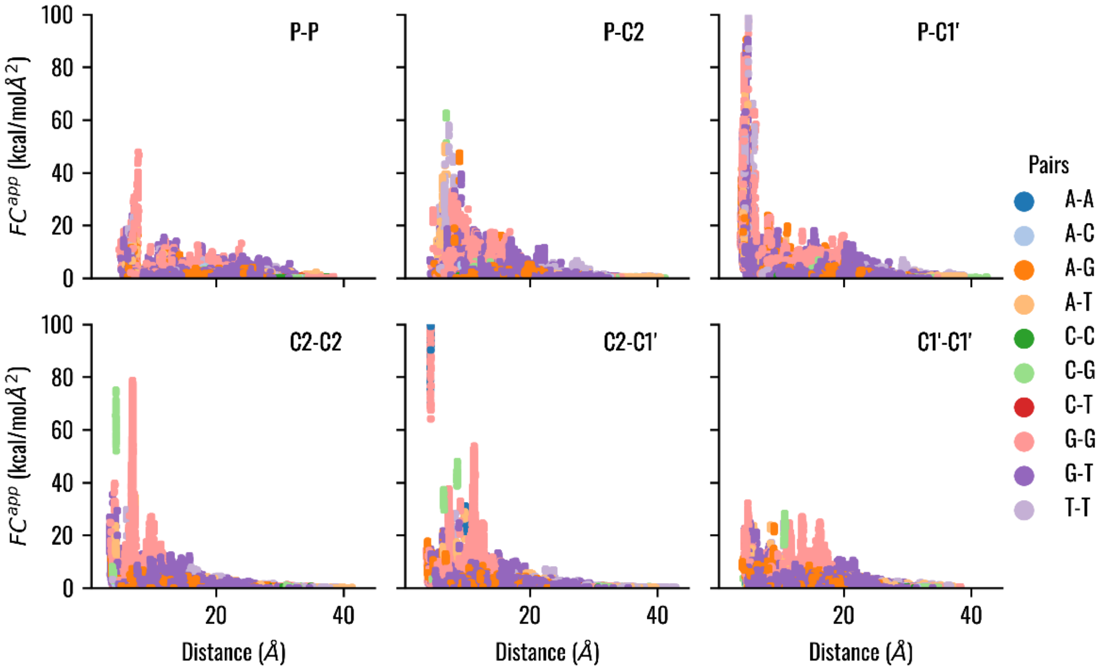


**Supplementary Figure 2.** Distribution of the apparent force constants $FC_{ij}^{app}$ as a function of the average distance $d_{ij}^{a}$ in the window in the training MD dataset of DNA-only systems. The apparent force constants between the different interacting couples are shown and the different nucleotide bases are highlighted with different colors.

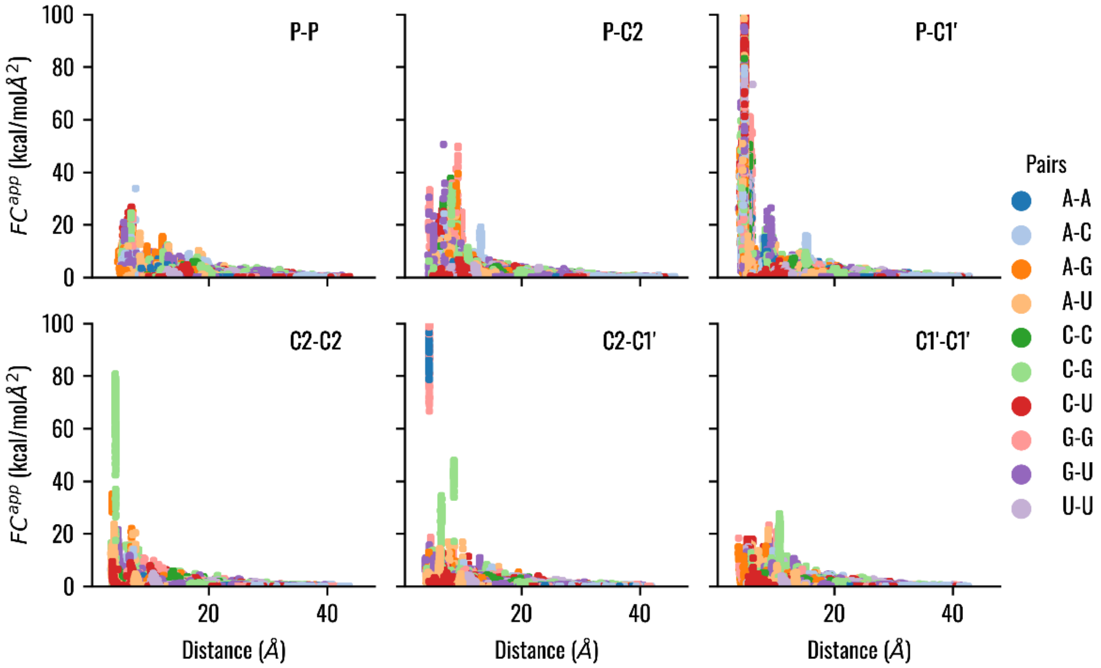


**Supplementary Figure 3.** Distribution of the apparent force constants $FC_{ij}^{app}$ as a function of the average distance $d_{ij}^{a}$ in the window in the training MD dataset of RNA-only systems. The apparent force constants between the different interacting couples are shown and the different nucleotide bases are highlighted with different colors.

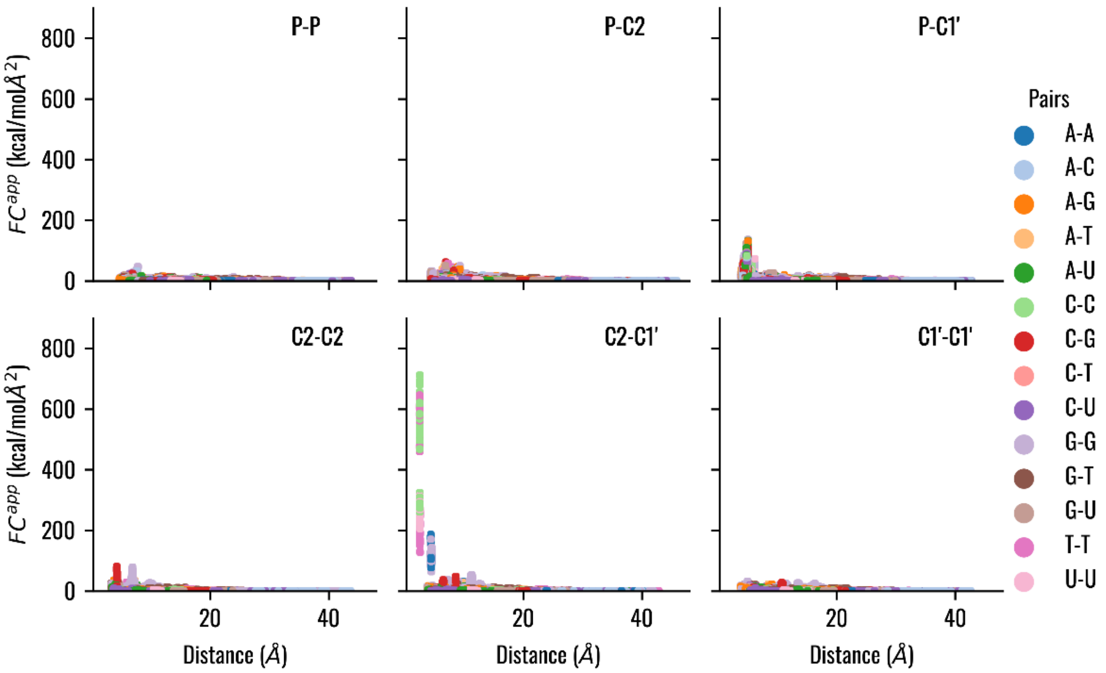


**Supplementary Figure 4.** Distribution of the apparent force constants $FC_{ij}^{app}$ as a function of the average distance $d_{ij}^{a}$ in the window in the training MD dataset of nucleic acids systems. The apparent force constants between the different interacting couples are shown and the different nucleotide bases are highlighted with different colors. The y-axis scale has been adapted to highlight the peaks in the C2-C1' contacts.

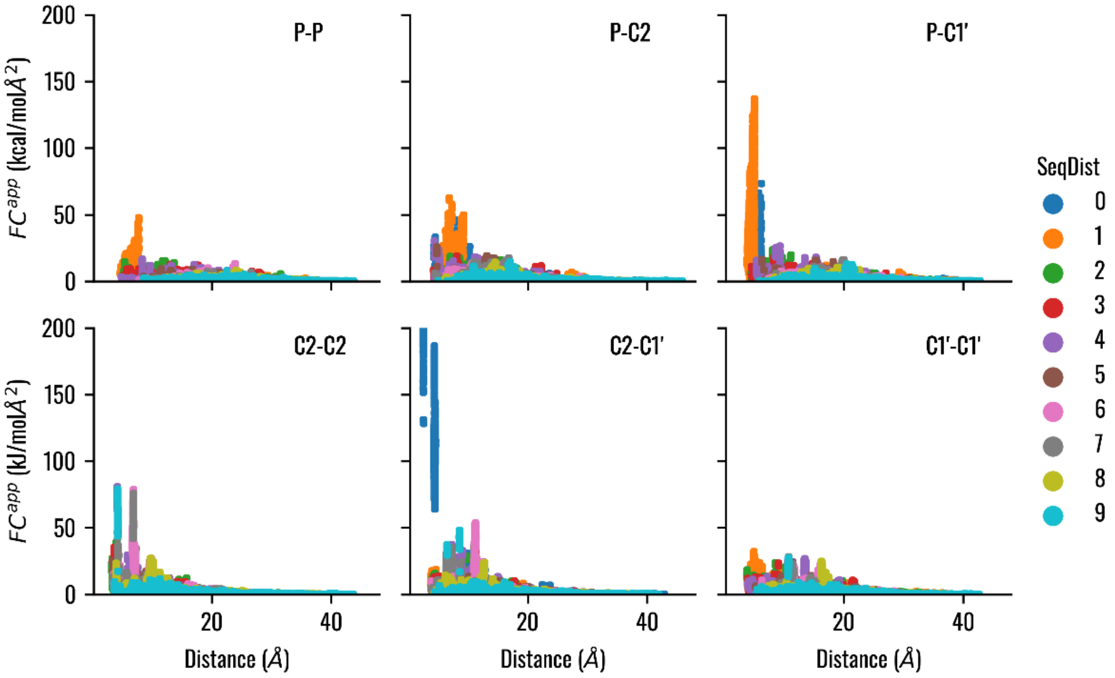


**Supplementary Figure 5.** Distribution of the apparent force constants $FC_{ij}^{app}$ as a function of the average distance $d_{ij}^{a}$ in the window in the training MD dataset of nucleic acids systems. The apparent force constants between the different interacting couples are shown and the sequential distance between the beads are highlighted with different colors.

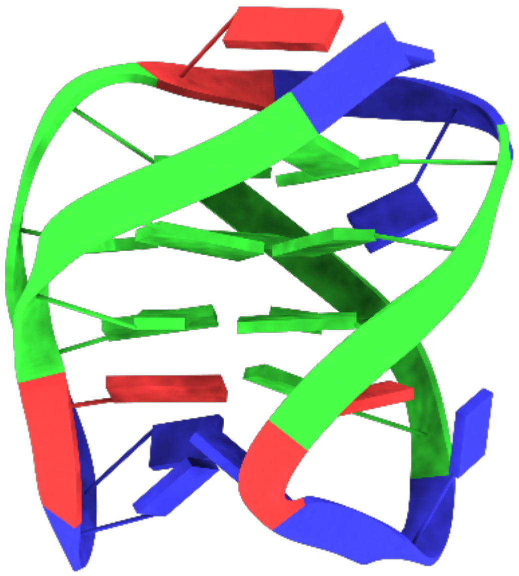

**Supplementary Figure 6.** Representation of a stable DNA β-hairpin with stacking interactions between guanines (green). The structure corresponds to the PDB 5M1W and has been simulated in Salsbury et al.[9].

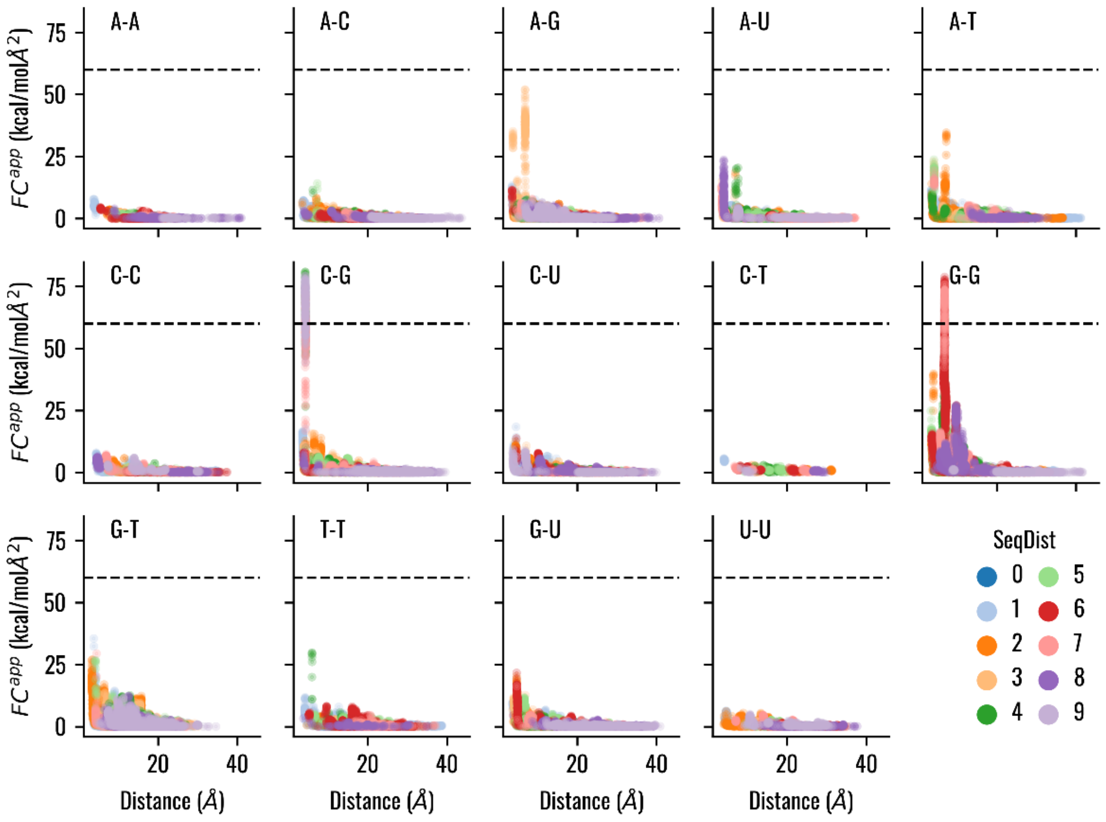


**Supplementary Figure 7.** Distribution of the C2-C2 apparent force constants $FC_{ij}^{app}$ as a function of the average distance $d_{ij}^{a}$ in the window in the training MD dataset of nucleic acids systems. The apparent force constants between the different interacting nucleotides are shown and the sequential distance between the beads are highlighted with different colors. Each point is semi-transparent, so darker regions correspond to higher density of data.

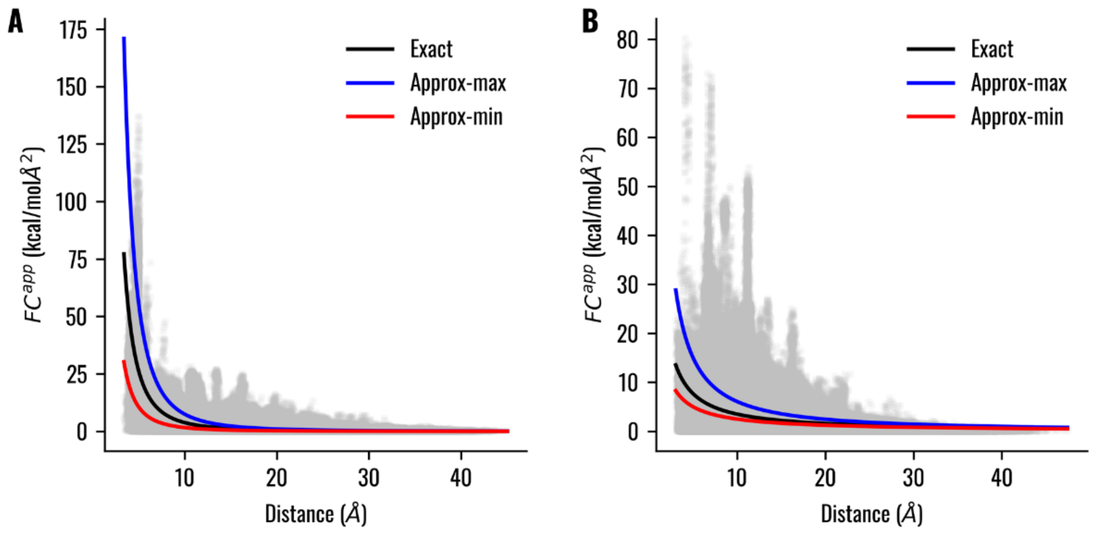


**Supplementary Figure 8.** Fitting intervals for the (A) "*pseudo-covalent*" and (B) "*Van der Waals*" interaction types. In the apparent force constant versus distance plot, the black line represents the function obtained by the fitting procedure, while the red and blue lines represent, respectively, the curves associated with minimum and maximum values of the parameters used in the random search optimization.

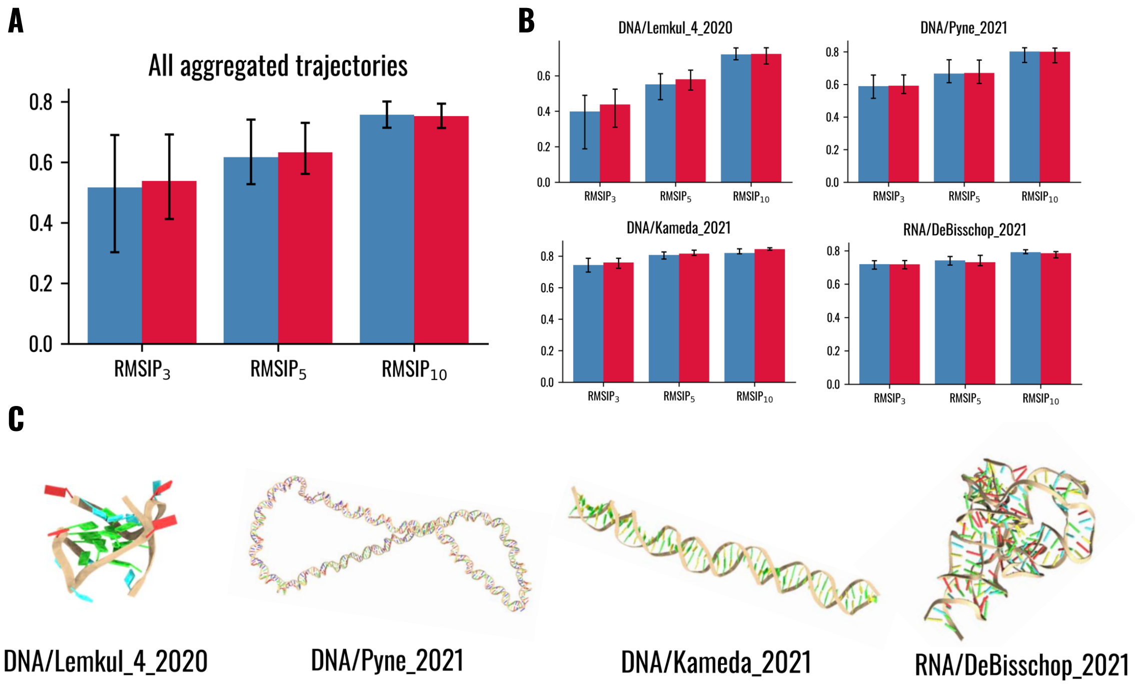


**Supplementary Figure 9.** Comparison between the uniform-spring ENM from Pinamonti et al.[26] (blue) and the edENM (red) in terms of agreement with the conformational dynamics observed in the four test MD simulations for DNA- and RNA-only systems. (A) Average *RMSIP* values ($RMSIP_3$, $RMSIP_5$, $RMSIP_{10}$) aggregated for the four testing MD simulations. Error bars refer to the first and third quantiles across the different MD trajectory windows. (B) Comparison with individual MD simulations from Lemkul[3], Pyne et al.[7], Kameda et al.[5], and De Bisschop et al.[15]. A graphical representation of each nucleic acid system is shown in (C).

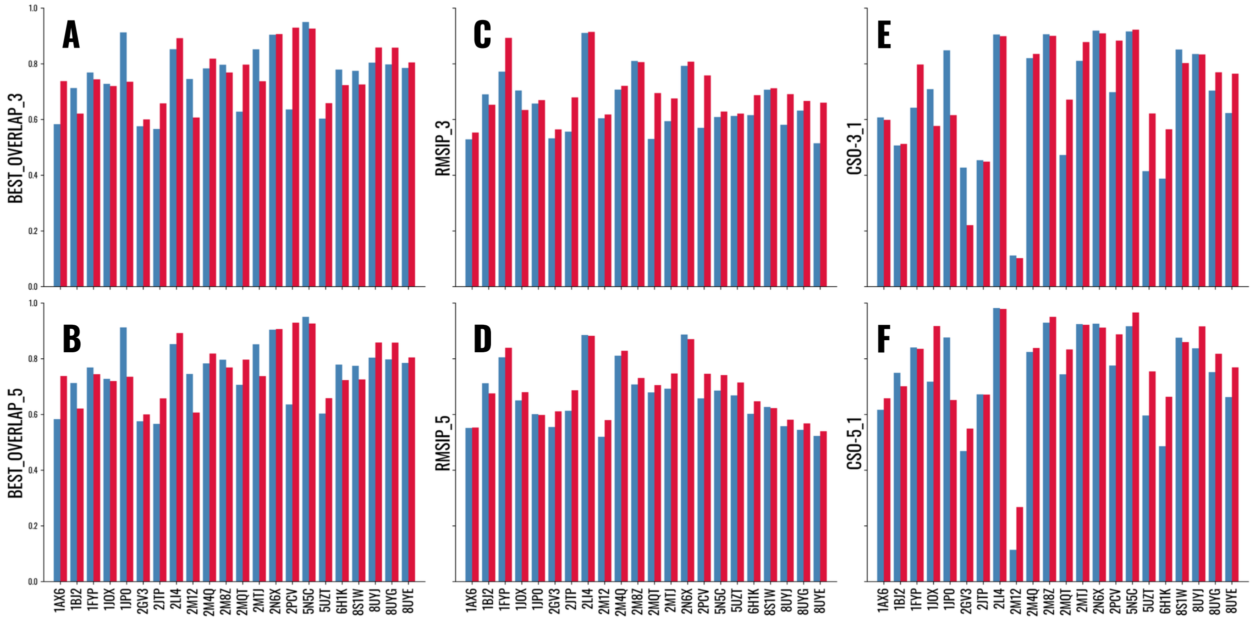


**Supplementary Figure 10.** Comparison between the uniform-spring ENM from Pinamonti et al.[26] (blue) and the edENM (red) and the experimental PCs from the test set of multi-model ensembles of DNA- and RNA-only systems from Supplementary Table 2: (A) $O_{max}$ for the first 3 sets of NM-PC modes; (B) $O_{max}$ for the first 5 sets of modes; (C) *RMSIP* for the first 3 sets of modes; (D) *RMSIP* for the first 5 sets of modes; (E) *CSO* for the first 3 NMs to cover PC1; (F) *CSO* for the first 5 NMs to cover PC1.

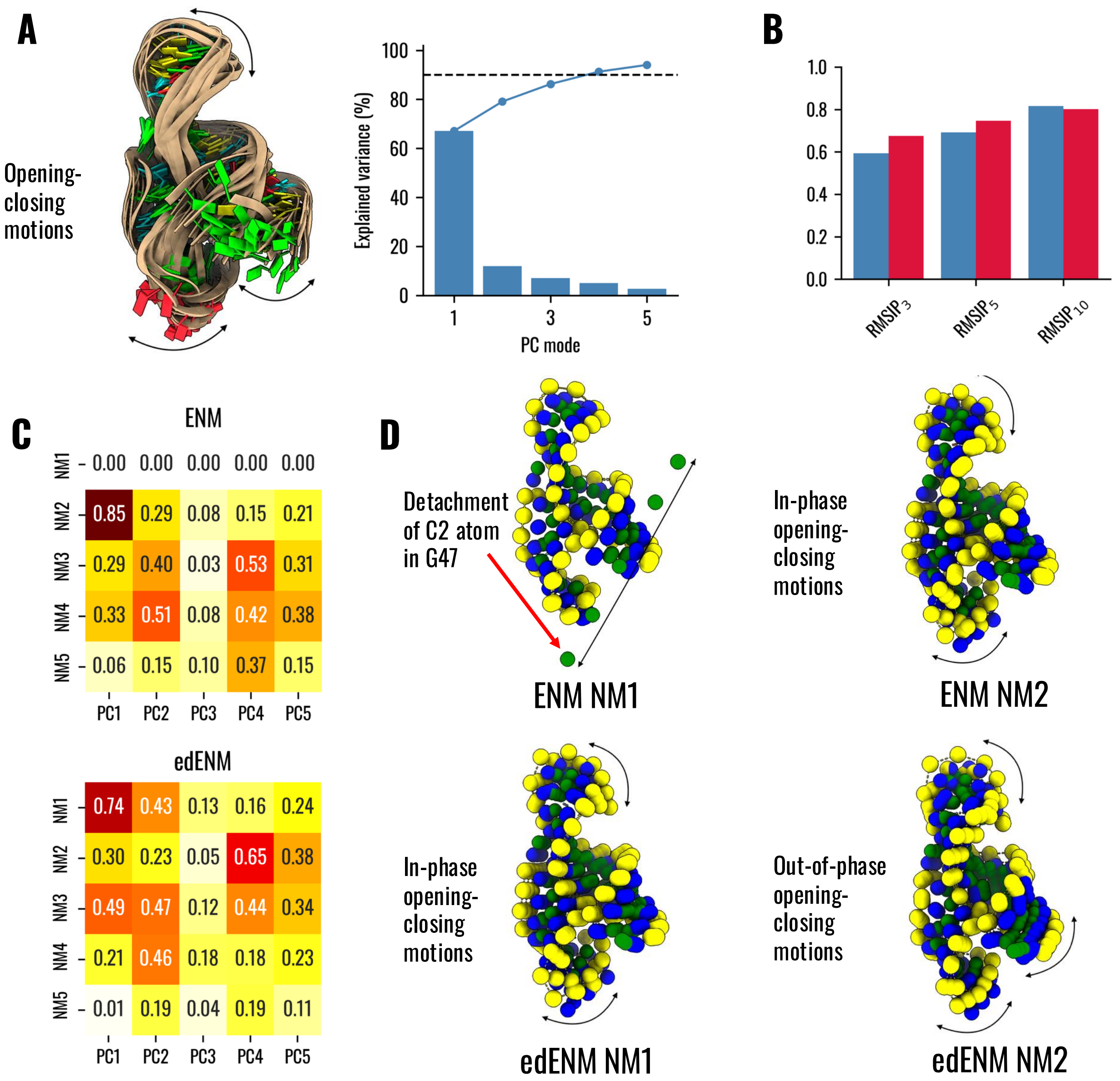


**Supplementary Figure 11.** Comparison between ENM, edENM and PC modes from the NMR ensemble (PDB: 2MTJ) of the III-IV-V three-way junction of VS ribozyme. (A) Ensemble of 21 models in the solution NMR ensemble used for PCA. The black arrows highlight the opening-closing motions apparent from the experimental ensemble, which dominates PC1. The graph shows the variance captured by each PC mode (blue bars), the cumulative variance of the first PC modes (continuous blue line), and a threshold of 90% variance (dashed black line). PC1 captures ~67% variance and PC1+PC2 together allow to capture ~80% variance in the ensemble. (B) Comparison between ENM (blue bars) and edENM (red) in terms of *RMSIP* scores (considering both 3, 5, and 10 modes) between NMs and experimental PCs. (C) Matrix of overlaps between the first 5 PC modes and the first 5 NMs from ENM and edENM. Maximum overlaps are found between NM2 and PC1 for ENM (0.85 overlap) and between NM1 and PC1 for edENM (0.74). (D) Graphical representation of the motions associated to NM1-NM2 for ENM (above) and edENM (below). Beads represent P, C1' and C2 positions and follow the same coloring as in Fig. 3. Note how both models allow to capture the opening-closing motions observed in the NMR ensemble, with NM2 of ENM with a ~11% better overlap than NM1 of edENM, but the main mode of ENM exhibits a detachment of the C2 bead in the terminal nucleotide G47, which makes NM1 completely uncorrelated to any experimental motion. This detachment is clearly prevented in the edENM due to stronger short-range interactions compared to the uniform-spring ENM.

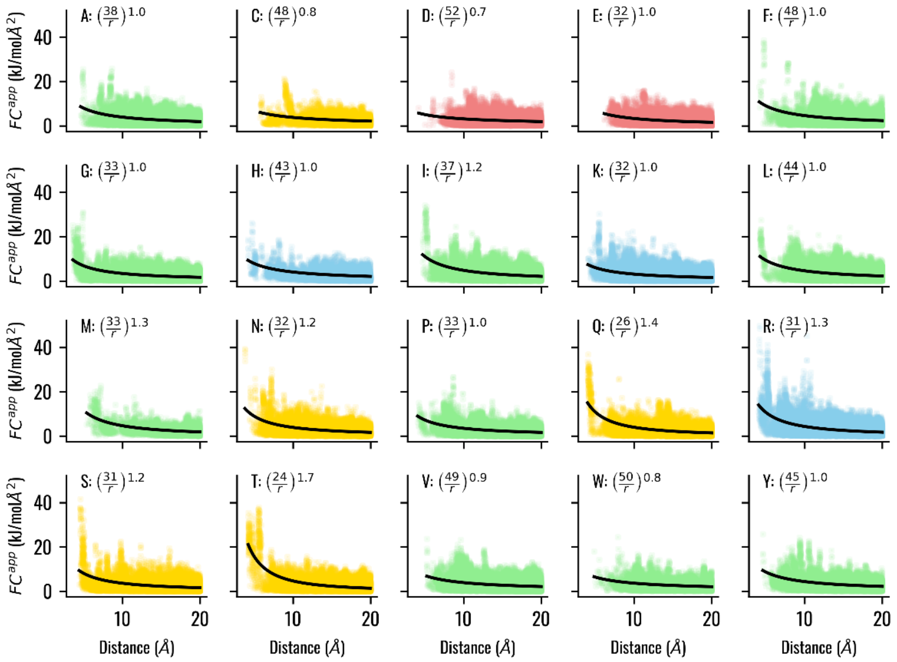

**Supplementary Figure 12.** Distribution of the C$^{\alpha}$-P apparent force constants $FC_{ij}^{app}$ as a function of the average distance $d_{ij}^{a}$ in the window in the training MD dataset of the protein-nucleic acid systems. The apparent force constants with the different amino acids are shown and the points are colored according to the amino acid type, with acidic, basic, polar, and hydrophobic residues highlighted in red, blue, yellow, and green, respectively. Each point is semi-transparent, so darker regions correspond to higher density of data. The black line indicates the function that best fits the data, whose mathematical expression is reported in the plot.

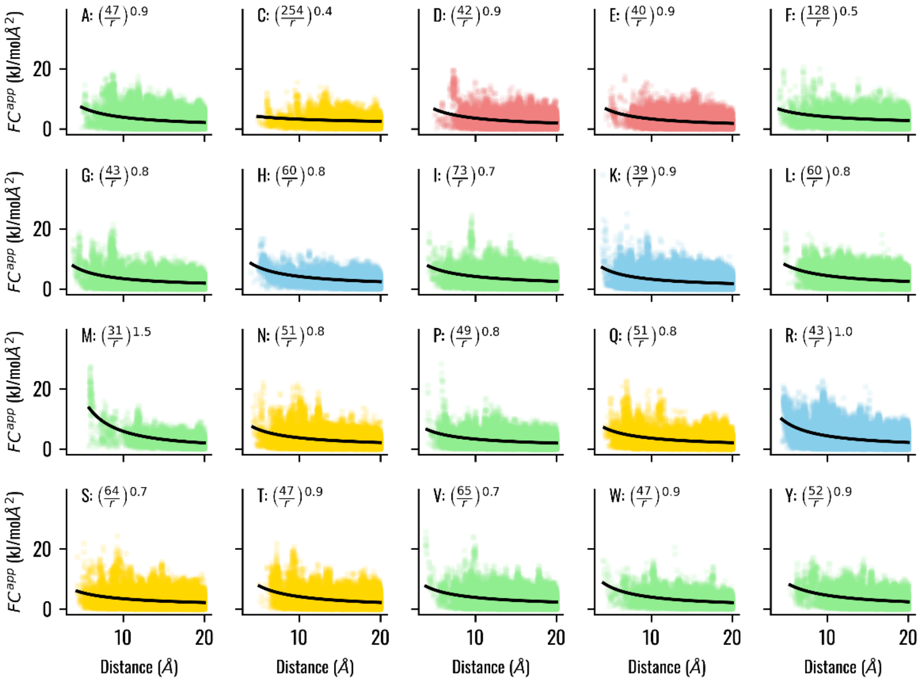

**Supplementary Figure 13.** Distribution of the C$^{\alpha}$-C1' apparent force constants $FC_{ij}^{app}$ as a function of the average distance $d_{ij}^{a}$ in the window in the training MD dataset of the protein-nucleic acid systems. The apparent force constants with the different amino acids are shown and the points are colored according to the amino acid type, with acidic, basic, polar, and hydrophobic residues highlighted in red, blue, yellow, and green, respectively. Each point is semi-transparent, so darker regions correspond to higher density of data. The black line indicates the function that best fits the data, whose mathematical expression is reported in the plot.

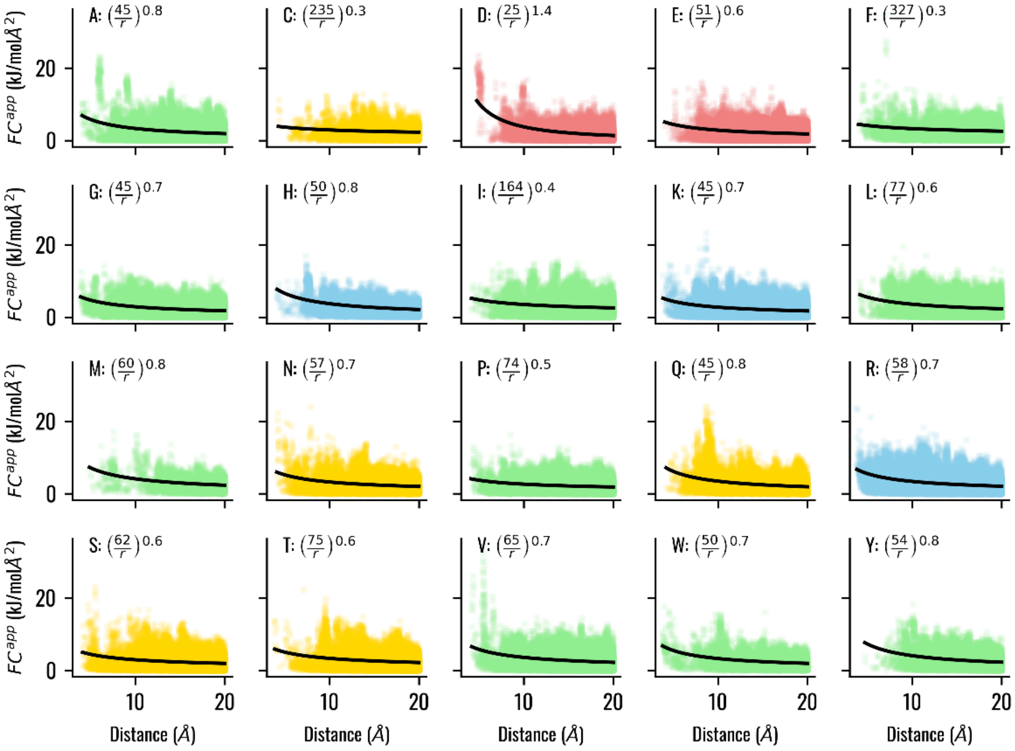


**Supplementary Figure 14.** Distribution of the $C^{\alpha}$-C2 apparent force constants $FC_{ij}^{app}$ as a function of the average distance $d_{ij}^{a}$ in the window in the training MD dataset of the protein-nucleic acid systems. The apparent force constants with the different amino acids are shown and the points are colored according to the amino acid type, with acidic, basic, polar, and hydrophobic residues highlighted in red, blue, yellow, and green, respectively. Each point is semi-transparent, so darker regions correspond to higher density of data. The black line indicates the function that best fits the data, whose mathematical expression is reported in the plot.

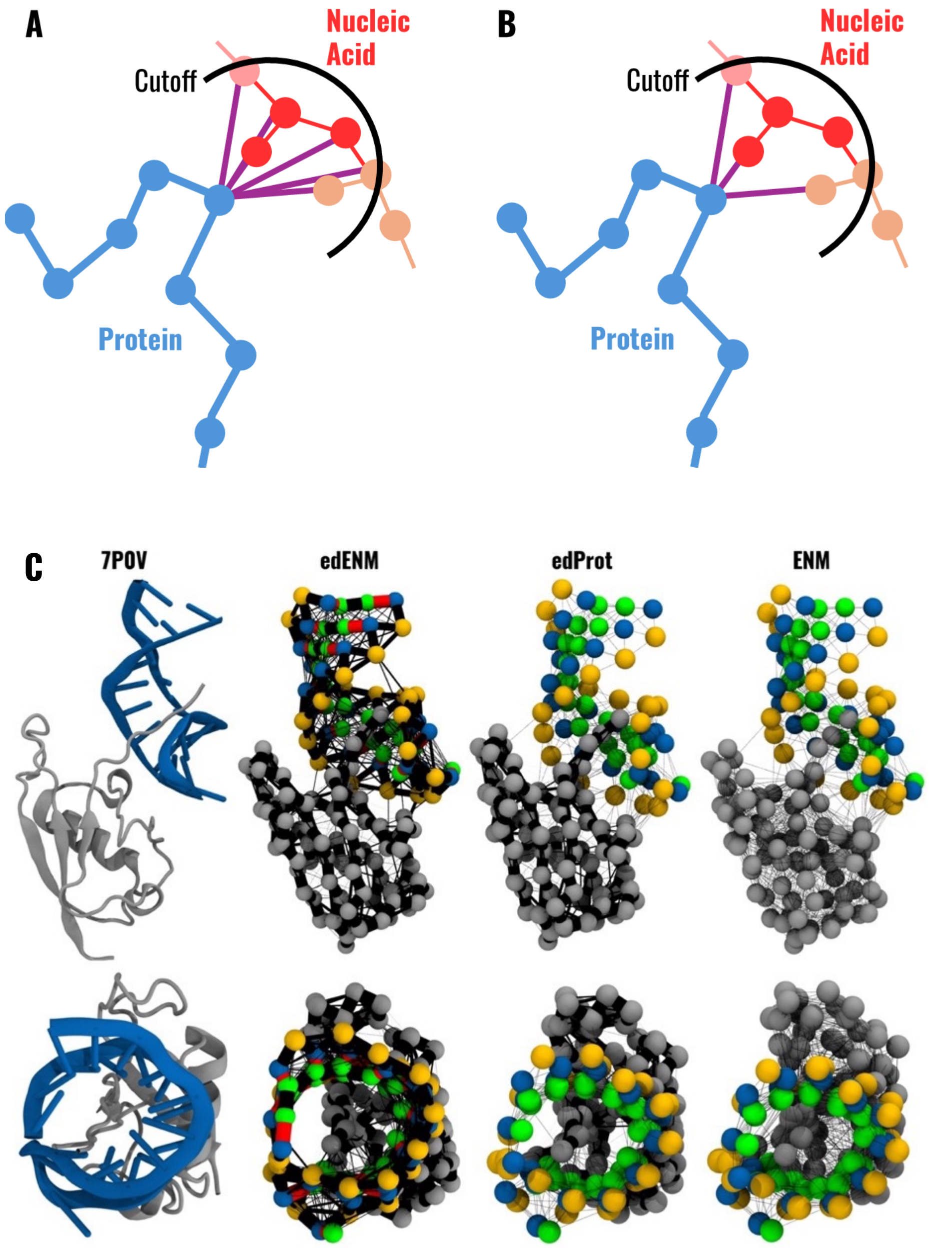


**Supplementary Figure 15.** (A,B): Representation of the (A) standard and (B) proposed approach to define the protein-nucleic acid contacts at the interface between the protein and the nucleic acid. In the standard approach, each protein bead is connected to all nucleic acid beads within a spatial cutoff, while in the proposed approach protein beads are only connected to one bead per nucleic acid residue, i.e., the bead that is closest to the protein $C^{\alpha}$ atom. In the figure, protein residues are shown as blue beads, and different residues in the nucleic acid with different colors (pink, red, orange), and protein-nucleic acid contacts as purple lines. The black spherical curve represents the space delimited by the spatial cutoff centered on each $C^{\alpha}$ atom. (C) Representation of different protein-nucleic acid network models for human SF3A1 ubiquitin-like domain in complex with U1 snRNA stem-loop 4 (PDB ID: 7P0V). The upper panel shows a lateral view, the lower panel a view from above. edENM represents our optimized network model for the whole protein-nucleic acid complex. edProt uses the edENM parametrization for the protein, but the standard ENM from Pinamonti et al.[26] for the nucleic acid. ENM uses uniform-spring ENM topologies for both. Bead colors and spring diameters are equivalent to those from Fig. 3 in the main text.

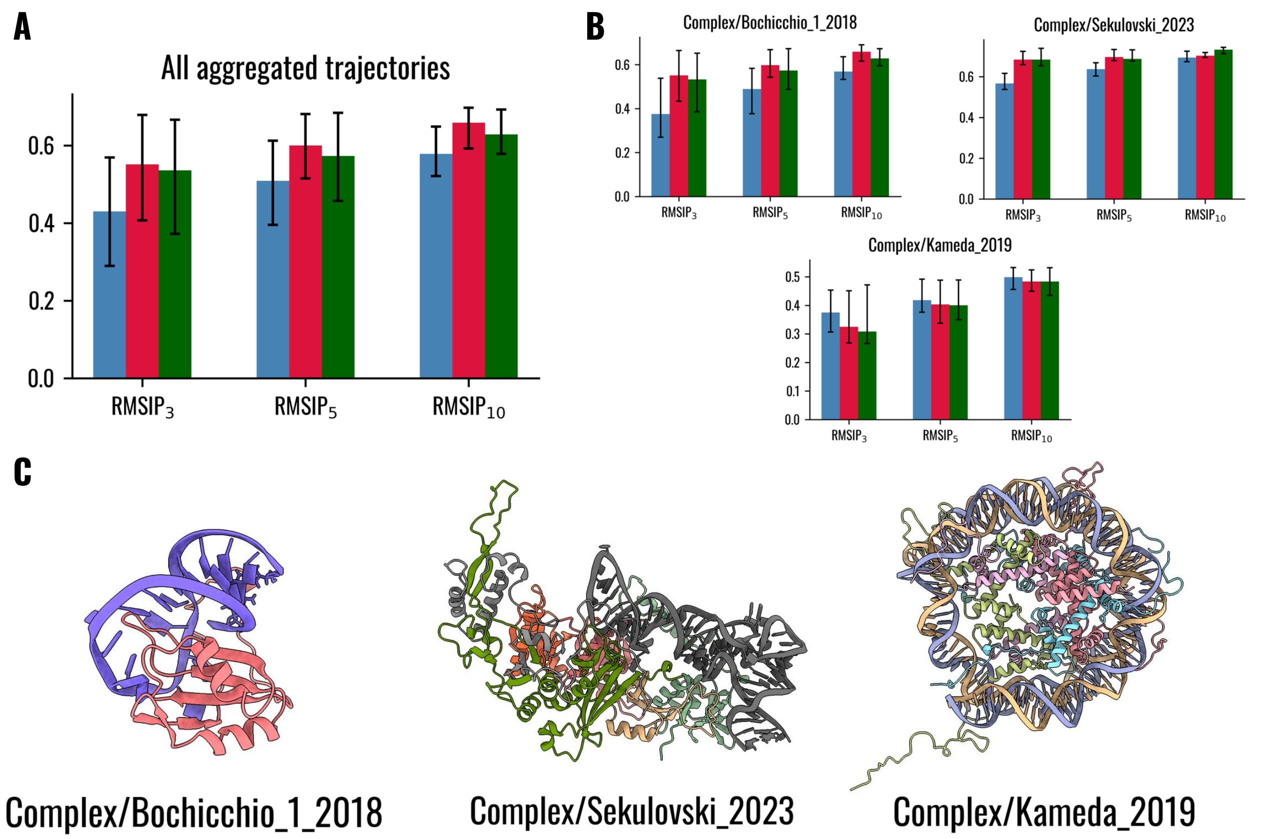


**Supplementary Figure 16.** Comparison between ENM (blue), edENM (red), and edProt (green) in terms of agreement with the conformational dynamics observed in the three test MD simulations for protein-nucleic acid systems. (A) Average *RMSIP* values ($RMSIP_3$, $RMSIP_5$, $RMSIP_{10}$) aggregated for the three testing MD simulations. Error bars refer to the first and third quantiles across the different MD trajectory windows. (B) Comparison with individual MD simulations from Bochicchio et al.[10], Sekulovski et al.[22], and Kameda et al.[17]. A graphical representation of each molecular system is shown in (C).

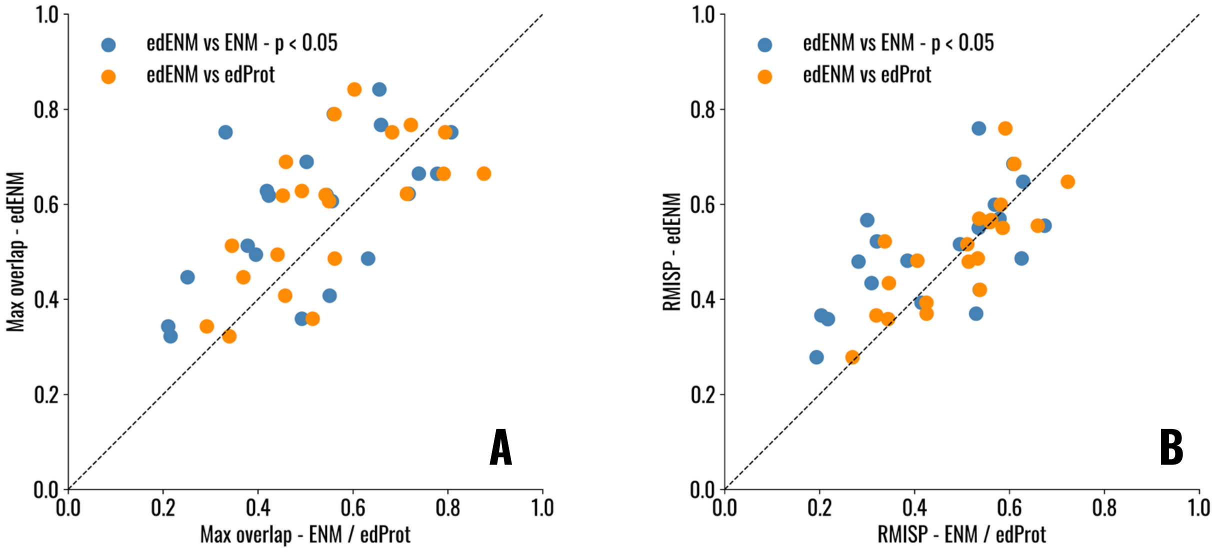


**Supplementary Figure 17.** Scatterplot showing the comparison between the edENM and ENM (blue points) or edProt (orange points) in terms of ability to capture experimental PCs from the set of NMR structures of protein-nucleic acid systems (Supplementary Table 3). (A) $O_{max}$ for the set of first 3 NMs and experimental PCs. (B) *RMSIP* for the set of first 3 NMs and experimental PCs. *P*-values arising from one-sided Wilcoxon tests between $O_{max}$ and *RMSIP* scores are reported in Supplementary Table 9.

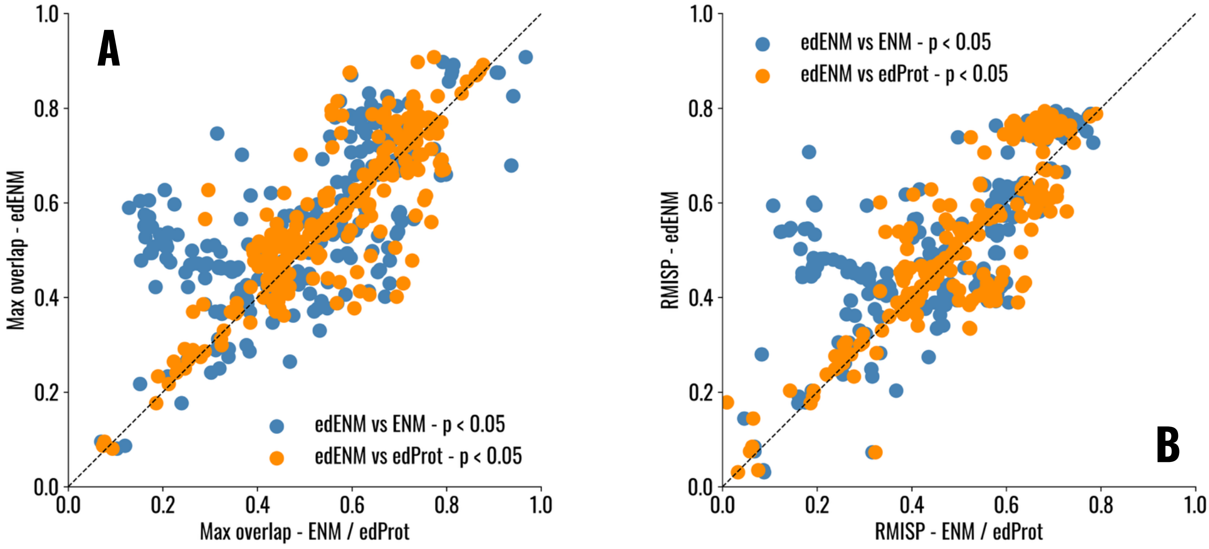


**Supplementary Figure 18.** Scatterplot showing the comparison between the edENM and ENM (blue points) or edProt (orange points) in terms of ability to capture experimental PCs from the set of protein-nucleic acid conformational ensembles at the level of individual PDB models (Supplementary Table 4). (A) $O_{max}$ for the set of first 3 NMs and experimental PCs. (B) *RMSIP* for the set of first 3 NMs and experimental PCs. Each point corresponds to an individual PDB. *P*-values arising from one-sided Wilcoxon tests between $O_{max}$ and *RMSIP* scores are reported in Supplementary Table 11.

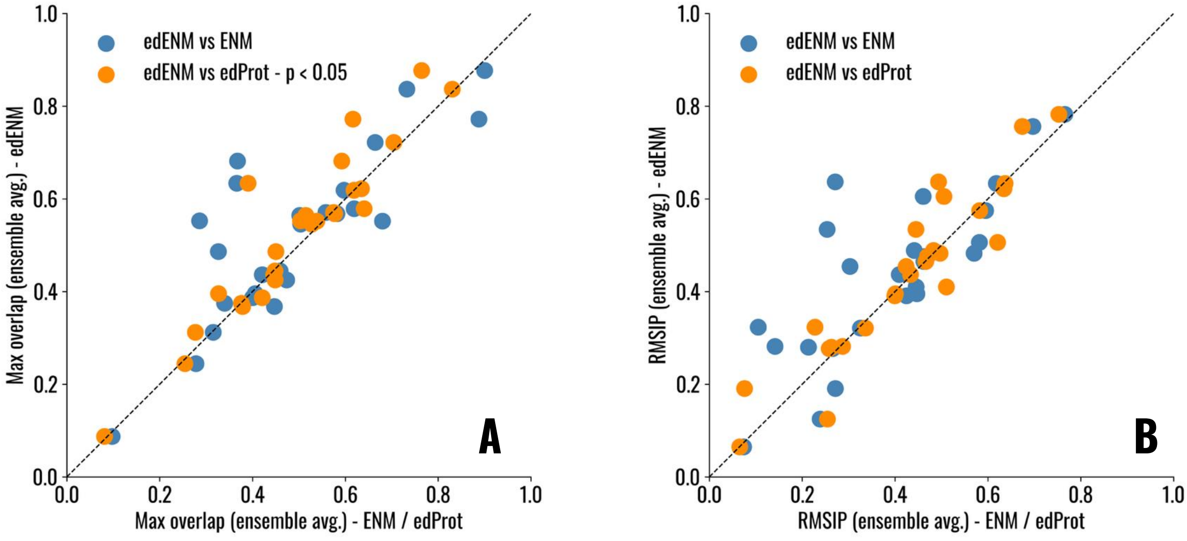


**Supplementary Figure 19.** Scatterplot showing the comparison between the edENM and ENM (blue points) or edProt (orange points) in terms of ability to capture experimental PCs from the set of protein-nucleic acid conformational ensembles, considering average similarity scores per ensemble. (A) $O_{max}$ for the set of first 3 NMs and experimental PCs. (B) *RMSIP* for the set of first 3 NMs and experimental PCs. Each point corresponds to the average score for each individual ensemble. *P*-values arising from one-sided Wilcoxon tests between $O_{max}$ and *RMSIP* scores are reported in Supplementary Table 12.

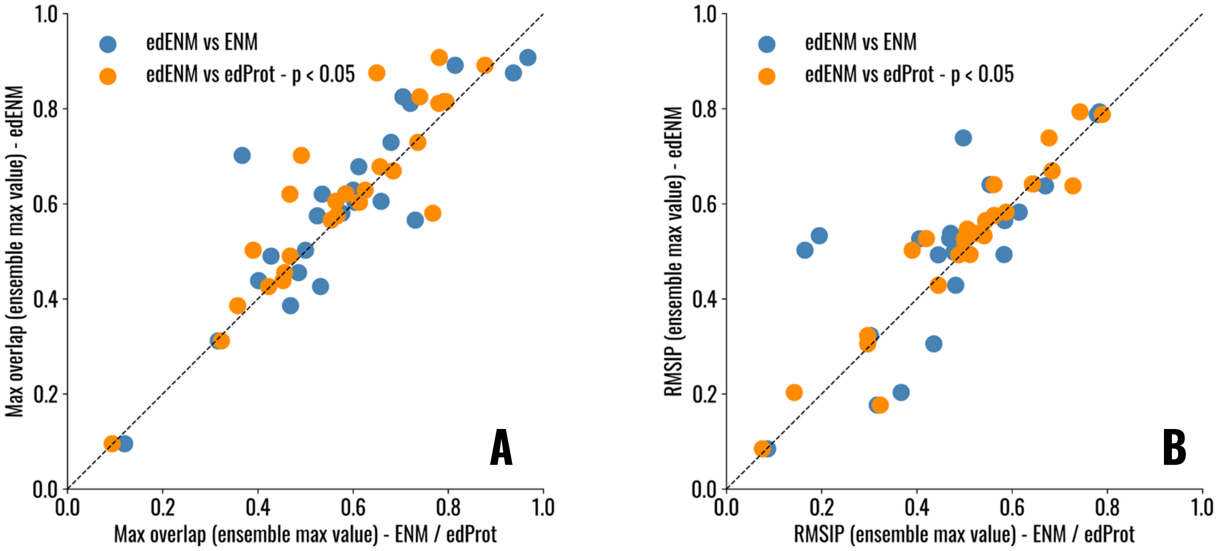


**Supplementary Figure 20.** Scatterplot showing the comparison between the edENM and ENM (blue points) or edProt (orange points) in terms of ability to capture experimental PCs from the set of protein-nucleic acid conformational ensembles, considering maximum similarity scores per ensemble. (A) $O_{max}$ for the set of first 3 NMs and experimental PCs. (B) *RMSIP* for the set of first 3 NMs and experimental PCs. Each point corresponds to the maximum score for each individual ensemble. *P*-values arising from one-sided Wilcoxon tests between $O_{max}$ and *RMSIP* scores are reported in Supplementary Table 13.

### References


1. Vreede, J., Pérez de Alba Ortíz, A., Bolhuis, P. G. & Swenson, D. W. H. Atomistic insight into the kinetic pathways for Watson–Crick to Hoogsteen transitions in DNA. *Nucleic Acids Research* **47**, 11069–11076 (2019).

2. Vreede, J. & Swenson, D. W. H. Molecular dynamics simulations of DNA. (2017).

3. Lemkul, J. A. Same fold, different properties: polarizable molecular dynamics simulations of telomeric and TERRA G-quadruplexes. *Nucleic Acids Research* **48**, 561–575 (2020).

4. Polêto, M., Michel, H. M., Ratnasinghe, B., Salsbury, A. M. & Lemkul, J. A. Nucleic Acid Dynamics. https://doi.org/10.17605/OSF.IO/2PFJ8 (2017) doi:10.17605/OSF.IO/2PFJ8.

5. Kameda, T., Suzuki, M. M., Awazu, A. & Togashi, Y. Structural dynamics of DNA depending on methylation pattern. *Phys. Rev. E* **103**, 012404 (2021).

6. Kameda, T., Suzuki, M. M., Awazu, A. & Togashi, Y. Structural dynamics of DNA depending on methylation pattern: Simulation dataset. Zenodo https://doi.org/10.5281/ZENODO.3992686 (2020).

7. Pyne, A. L. B. *et al.* Base-pair resolution analysis of the effect of supercoiling on DNA flexibility and major groove recognition by triplex-forming oligonucleotides. *Nat Commun* **12**, 1053 (2021).

8. Pyne, A. L. B. *et al.* Base-pair resolution analysis of the effect of supercoiling on DNA flexibility and major groove recognition by triplex-forming oligonucleotides. (2021).

9. Salsbury, A. M., Michel, H. M. & Lemkul, J. A. Ion-Dependent Conformational Plasticity of Telomeric G-Hairpins and G-Quadruplexes. *ACS Omega* **7**, 23368–23379 (2022).

10. Bochicchio, A. *et al.* Molecular basis for the increased affinity of an RNA recognition motif with re-engineered specificity: A molecular dynamics and enhanced sampling simulations study. *PLoS Comput Biol* **14**, e1006642 (2018).

11. Bochicchio, A. *et al.* Molecular Basis For The Increased Affinity Of An Rna Recognition Motif With Re-Engineered Specificity: A Molecular Dynamics And Enhanced Sampling Simulations Study- Part 2. Zenodo https://doi.org/10.5281/ZENODO.1310053 (2018).

12. Bochicchio, A. *et al.* Molecular Basis For The Increased Affinity Of An Rna Recognition Motif With Re-Engineered Specificity: A Molecular Dynamics And Enhanced Sampling Simulations Study.-Part 8. Zenodo https://doi.org/10.5281/ZENODO.1465433 (2018).

13. Levintov, L. & Vashisth, H. Role of conformational heterogeneity in ligand recognition by viral RNA molecules. *Phys. Chem. Chem. Phys.* **23**, 11211–11223 (2021).

14. Levintov, L. & Vashisth, H. Dataset for: Role of conformational heterogeneity in ligand recognition by viral RNA molecules. Zenodo https://doi.org/10.5281/ZENODO.4521164 (2021).

15. De Bisschop, G. *et al.* Progress toward SHAPE Constrained Computational Prediction of Tertiary Interactions in RNA Structure. *ncRNA* **7**, 71 (2021).

16. De Bisschop, G. *et al.* Progress Toward SHAPE Constrained Computational Prediction of Tertiary Interactions in RNA Structure. Zenodo https://doi.org/10.5281/ZENODO.5642809 (2021).

17. Kameda, T., Awazu, A. & Togashi, Y. Histone Tail Dynamics in Partially Disassembled Nucleosomes During Chromatin Remodeling. *Front. Mol. Biosci.* **6**, 133 (2019).

18. Kameda, T., Awazu, A. & Togashi, Y. Histone tail dynamics in partially disassembled nucleosomes during chromatin remodeling: Simulation dataset. Zenodo https://doi.org/10.5281/ZENODO.2548539 (2019).

19. Baltrukevich, H. & Bartos, P. RNA-protein complexes and force field polarizability. *Front. Chem.* **11**, (2023).

20. Baltrukevich, H. & Bartos, P. Cas12j-RNA simulations in ff19SB+OL3, ff14SB+OL3, OPLS4 and AMOEBA force fields. Zenodo https://doi.org/10.5281/ZENODO.7694878 (2023).

21. Baltrukevich, H. & Bartos, P. RIG-I and RNA complex MD simulations in ff19SB+OL3, ff14SB+OL3, OPLS4 and AMOEBA force fields. Zenodo https://doi.org/10.5281/ZENODO.7695265 (2023).

22. Sekulovski, S., Sušac, L., Stelzl, L. S., Tampé, R. & Trowitzsch, S. Structural basis of substrate recognition by human tRNA splicing endonuclease TSEN. *Nat Struct Mol Biol* **30**, 834–840 (2023).

23. Trowitzsch, S. & Stelzl, L. Protein-RNA complex simulation. Zenodo https://doi.org/10.5281/ZENODO.6513519 (2022).

24. Riccardi, E., Van Mastbergen, E. C., Navarre, W. W. & Vreede, J. Predicting the mechanism and rate of H-NS binding to AT-rich DNA. *PLoS Comput Biol* **15**, e1006845 (2019).

25. Vreede, J. Predicting the mechanism and rate of H-NS binding to AT-rich DNA. (2019).

26. Pinamonti, G., Bottaro, S., Micheletti, C. & Bussi, G. Elastic network models for RNA: a comparative assessment with molecular dynamics and SHAPE experiments. *Nucleic Acids Res* **43**, 7260–7269 (2015).
